# Glutamate receptor-dependent cytosolic acidification in hippocampal neurons involves passive flux of protons from the extracellular space

**DOI:** 10.1101/2024.11.17.624027

**Authors:** Sofie E. Pedersen, Ainoa Konomi-Pilkati, Nikolaj W. Hansen, Trine Kvist, Leonie P. Posselt, Thorvald F. Andreassen, Andreas T. Sørensen, Jean François Perrier, Kenneth L. Madsen

## Abstract

Glutamate receptor-dependent cytosolic acidification can be induced in hippocampal neurons by pharmacological or seizure-like stimulation. This acidification is thought to arise from Ca^2+^ and metabolism-related processes, however, the exact underlying mechanism as well as its functional role remains uncertain. To reassess the mechanism of cytosolic acidification in excitatory hippocampal neurons and address the physiological relevance of the phenomenon, we combined pH/Ca^2+^ biosensors to study activity-induced pH dynamics in hippocampal neurons. First, we addressed cytosolic acidification in relation to LTP at hippocampal CA3-CA1 synapses. Using hippocampal slices from adult rats of both sexes, we show that LTP-inducing stimulation at the Schaffer collaterals evokes transient cytosolic acidification in hippocampal CA1 neurons. This highlights neuronal pH shifts as a trait of general hippocampal neurotransmission rather than a marker of excitotoxicity, possibly serving as a secondary messenger. Moreover, using dissociated hippocampal neurons from rat embryos, we show that glutamate receptor agonists typically induce larger cytosolic acid shifts compared to simple depolarization or spontaneous activity, suggesting that glutamate receptor-mediated acidification involves several separate mechanisms; pyruvate-dependent dampening of neuronal acidification may reflect a direct inhibition of NMDA receptors rather than reduced glycolytic activity, questioning the previously reported involvement of metabolism in cytosolic acidification; and whereas acid shifts induced by simple depolarization show exclusive dependence on cytosolic Ca^2+^, AMPA-induced acidification depends both on cytosolic Ca^2+^ *and* on an inward electrochemical driving force for protons. These results suggest that glutamate receptor-induced cytosolic acidification relies both on cytosolic Ca^2+^ and on a passive proton influx, possibly mediated by the receptor itself.

**Graphical abstract:** 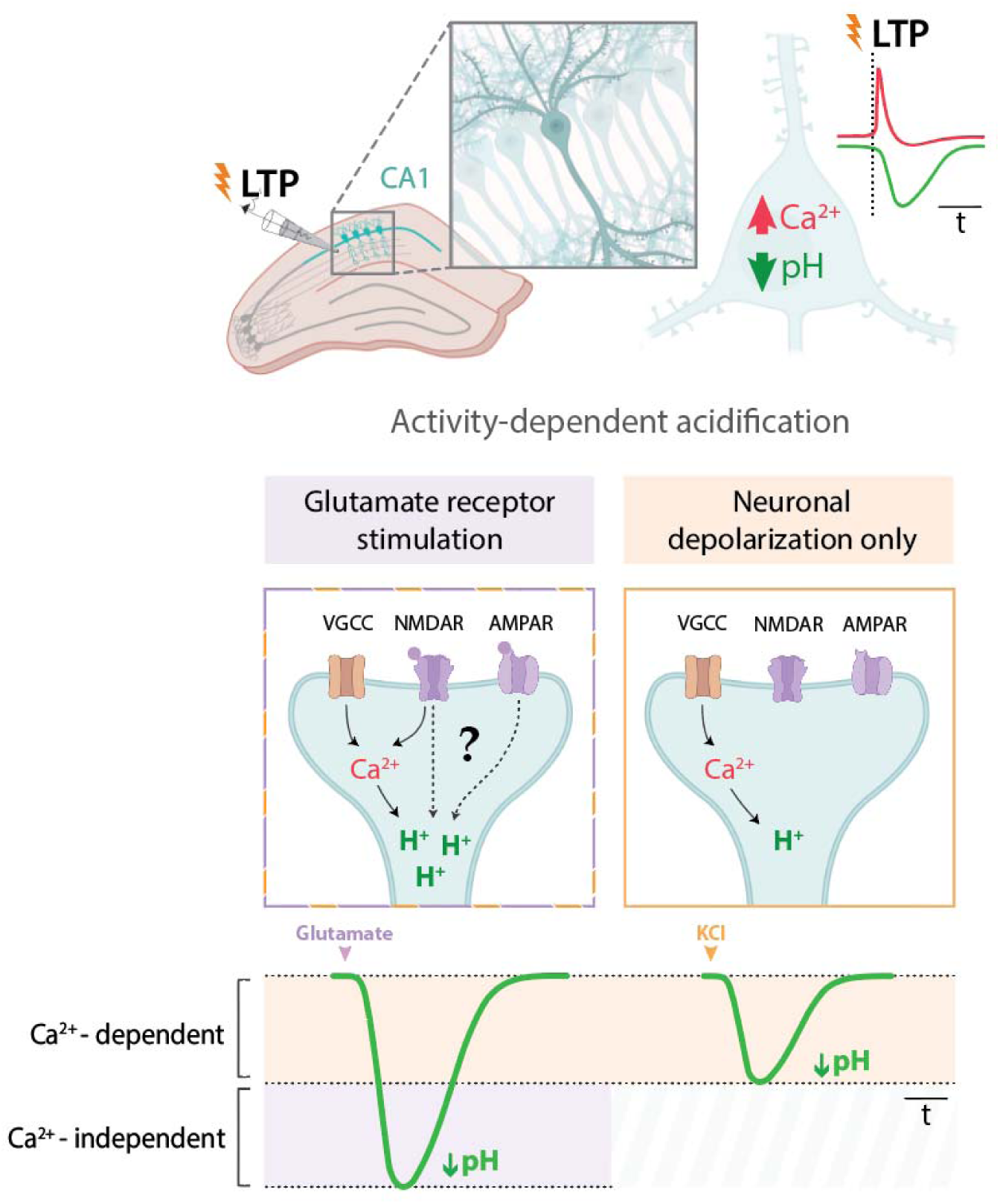

**Significance statement:** Although several studies report that hippocampal pyramidal neurons show significant cytosolic acidification in response to activation by drugs, epilepsy and stroke, the molecular mechanism and functional impact of the phenomenon remains uncertain. Using live imaging of both Ca and H dynamics, we demonstrate that the induction of LTP at hippocampal synapses is associated with cytosolic acidification in the postsynaptic neurons, suggesting that the cytosolic acid shifts may be a general trait of neurotransmission and related plasticity. We further revisit the mechanistic relation between the two ion systems. Our results suggest that glutamate receptor-induced cytosolic acidification involves two distinct mechanisms, one related to cytosolic Ca^2+^ and another involving a passive H influx, possibly mediated by the receptor itself.

## Introduction

Sustained pH homeostasis is essential for neuronal survival and function. Most proteins, including enzymes, ion channels and receptors, rely on pH to maintain their structure and function. (Tang et al. 1990, Traynelis et al., 1990, Billups et al., 1996, Krishek et al., 1996, Tombaugh et al., 1997). Moreover, intracellular pH gradients fuel vital cellular processes critical for neurotransmission, including mitochondrial electron-transport (driving ATP production), as well as loading and trafficking of neurotransmitter vesicles (Obara et al., 2008; Casey et al., 2010, Blakely et al., 2012). Consequently, alterations in neuronal pH have the potential to modulate neuronal activity and neurotransmission (Lee et al., 1996).

In hippocampal neurons and other nerve cells, excitatory synaptic activity not only increases cytosolic Ca^2+^ levels—a known mechanism of neuronal signaling—but also induces transient intracellular acidification (Hartley and Dubinsky 1993, Irwin et al. 1994, Koch and Barish 1994, Wang et al., 1994, Raimondo et al., 2012). While Ca^2+^ transients are a well-established marker of neuronal activity (Grienberger and Konnerth, 2012), the functional implications of activity-induced pH shifts remain less understood. These pH fluctuations have been observed during seizure-like activity (Siesjö et al., 1985; Xiong et al., 2000; Raimondo et al., 2012) and in response to pharmacological glutamate receptor activation or membrane depolarization (Hartley and Dubinsky, 1993; Irwin et al., 1994; Zhan et al., 1998; Svichar et al., 2011; Rathje et al., 2013).

The neuronal pH shifts associated with excitatory activity are thought to be generated from both Ca²⁺-dependent and metabolic processes (for a review see (Chesler, 2003)). In line with this, the absence of extracellular and cytosolic Ca^2+^ is consistently reported to reduce pharmacologically induced cytosolic acidification, see for example (Irwin et al., 1994; Wang et al., 1994; Zhan et al., 1998; Cheng et al., 2008). The underlying mechanism of this Ca²⁺-dependence is not well-established, but it has both been proposed to involve mitochondrial Ca^2+^ overload (Wang et al., 1994) and Ca²⁺-H^+^ exchange by the plasma membrane Ca²⁺-ATPase (PMCA) (Cheng et al., 2008). Activity-induced acceleration of neuronal metabolism has likewise been proposed to produce cytosolic acidification. Glycolysis, mitochondrial respiration, and the astrocyte-neuron-lactate-shuttle (ANLS) are all processes known to generate acids, which may be released into the cytosol and create acidification (Casey et al., 2010). Accordingly, a few studies have identified glycolysis as the main contributor to pharmacologically induced neuronal acidification (Zhan et al., 1997)(Zhan et al., 1998).

In the present study, we employ a dual biosensor approach (co-expression of red-shifted Ca^2+^ sensor (R-GECO) and green fluorescent pH sensor (cyto-pHluorin)) to characterize the relationship between cytosolic Ca^2+^ and pH, dual-expression of red-shifted Ca^2+^ sensor (RGECO) and green pH sensor (cyto-pHluorin), in single neurons. We examine the fluorescent changes from neurons responding to synaptic stimulation or pharmacological activation and re-assess the proposed mechanisms behind activity-induced acid shifts.

By applying the dual biosensor approach in hippocampal slices, we show that cytosolic acidification is induced in CA1 neurons in response to a conventional LTP-inducing theta-burst stimulation applied at the Schaffer collaterals, alongside transient Ca^2+^ increase. These results suggest that the cytosolic acidification in CA1 neurons, induced by excitatory synaptic stimulation, have both Ca²⁺- and H^+^-gradient dependent components, and that the phenomenon may be a trait of physiological neurotransmission at CA3-CA1 synapses. Moreover, using dissociated hippocampal cultures, we show that NMDA and AMPA receptor activation typically generates larger acid shifts compared to membrane depolarization alone. Our data also suggest that pyruvate can directly reduce NMDA receptor efficacy, which questions the previously proposed main role for glycolysis in glutamate-induced acidification. Additionally, we show that pharmacologically induced acid shifts are strongly dependent on the presence of extracellular Ca^2+^, but that a cytosolic Ca^2+^ rise *per se* is not sufficient to induce acid shifts of comparable magnitudes to NMDA and AMPA responses. Finally, we reveal that AMPA-induced acidification in part relies on an inward plasma membrane H^+^ gradient, which is not the case for acidification caused by simple depolarization. Together these results indicate that activation of glutamate receptors allows for an influx of protons, putatively through the receptors themselves.

## Materials and Methods

### Chemicals

Unless otherwise stated, all buffer reagents were purchased from Sigma-Aldrich (St. Louis, MO).

### Adeno-associated virus constructs and stereotaxic injections

Ready-to-use adeno-associated virus (AAV) particles encoding synapsin-driven Ca^2+^ sensor, jRGECO1a, were purchased from Addgene (pAAV9.Syn.NES-jRGECO1a.WPRE.SV40 from Douglas Kim and GENIE project, Addgene plasmid #100854)(Dana et al., 2016). Recombinant AAV9 vector encoding pH-sensor, cyto-pHluorin, was designed as follows: a 1289 bp mouse CamKII promoter flanking the 5’-ITR site was placed upstream of the coding sequence of super-ecliptic pHluorin, and terminated by WPRE and hGH poly(A) signal at the 3’-ITR end. All cloning steps were performed with traditional PCR cloning techniques, and the vector was sequenced for verification of correct configuration and integration. High-titer AAV particles (~2.6 × 10^13^ genome copies per mL) were produced in-house according to procedures described elsewhere (Sørensen et al., 2016). For use, jRGECO1a and cyto-pHluorin AAVs were mixed and diluted with sterile Dulbecco’s phosphate-buffered saline, achieving final titers of ~2.5×10^12^ genome copies per mL.

AAV infusions were carried out by stereotaxic injections in four weeks old C57bl6N mice of mixed gender (Janvier Lab, Le Genest-Saint-Isle, France) under general isoflurane anesthesia (2% during induction, 1.5–2% during maintenance) in a stereotaxic instrument (Kopf Instruments, Tujunga, CA, USA). A midline incision was made down the scalp and a craniotomy was made for bilateral infusions into the two different areas within the hippocampalCA1 region. The AAV mix was infused with two depositions of 300 nL per infusion site, using the following pre-set coordinates - *Left side:* from bregma: AP −2.70 mm, ML −2.89 mm, from skull: DV 2.25 and 2.10 mm; *Right side:* AP −2.10 mm, ML +1.80 mm, from skull: DV 1.50 and 1.30 mm. The pre-set infusion coordinates, based on the average brain size of C57BL/6J mice (Franklin and Paxinos, 2008), were adjusted to the smaller sized brains of the young animals, using the bregma-lambda distance as a correction factor, see (Moore and Boehm, 2009). Analgesic and antibiotic mixture (Norodyl: 0.5 mg/mL, Baytril: 0.5 mg/mL) was administered subcutaneously before and until two days after surgery. The animals were used in experiments three-four weeks after AVV injections.

### Brain slice preparation

Sagittal slices (300 µm) from AAV injected mice (7-8 weeks old) containing the hippocampus were prepared in ice-cold, carbogen (95% O_2_/5% CO_2_) saturated sucrose substituted artificial cerebrospinal fluid (sACSF) containing (in mM): Sucrose (200), NaHCO_3_ (25), glucose (11), KCl (3), CaCl_2_ (0.1), MgCl_2_ (4) and KH_2_PO_4_ (1.1), Sodium pyruvate (2), myoinositol (3) and ascorbic acid (0.5) using a vibratome (Leica VT1200, Germany). Brains slices were transferred to an interface type holding chamber filled with 28 °C carbogen saturated ASCF containing (in mM): NaCl (111), NaHCO_3_ (25), glucose (11), KCl (3), CaCl_2_ (2.5), MgCl_2_ (1.3) and KH_2_PO_4_ (1.1) and allowed to recover for at least 1½ hour.

### Combined electrophysiology and two-photon fluorescence imaging

Brain slices were placed in a submerged type recording chamber under a 16X objective of a combined two-photon and electrophysiology microscope (Bergamo II; Thorlabs, USA). The slices were continuously perfused with carbogen-saturated aCSF at room temperature. A bipolar tungsten stimulation electrode (TM33CCNON; World Precision Instruments, Sarasota, FL, USA) connected to a stimulus isolator (WPI A365, USA) was positioned in the stratum radiatum region at the CA3-CA1 border in the dorsal hippocampus. Field EPSPs (fEPSPs) were evoked by delivering 0.05 ms current pulses of 250-370 µA to Schaffer collaterals. Evoked fEPSPs were recorded from the stratum radiatum of CA1 using aCSF filled extracellular glass electrodes (4-7 MΩ), pulled on a horizontal puller (Sutter P-87, USA), connected to a Multiclamp 700B amplifier (Molecular Devices, USA) via a CV-7B Head stage. Signals were sampled at 10 KHz using a 1440A digitizer (Molecular Devices, USA) and displayed on a computer using Clampex software (Molecular Devices, USA). After establishing a stable baseline for fEPSP, long term potentiation (LTP) was induced with a single theta-burst protocol consisting of 20 bursts of 4 stimulations at 100 Hz repeated every 200 ms.

Simultaneous Ca²⁺- and pH imaging of CA1 pyramidal cells co-expressing jRGECO1a and cyto-pHluorin was performed before, during and after LTP induction by means of two-photon microscopy. jRGECO1a and cyto-pHluorin sensors were excited at a wavelength of 980 nm using a Ti:Sapphire laser (Coherent Chameleon Ultra 2, USA), and the emitted fluorescence was collected by two PMTs (BDM3214; Thorlabs; USA) using a 525/50 nm BrightLine® single-band bandpass and 607/70 nm BrightLine® single-band bandpass filters (Semrock; USA). Signal were acquired using ThorImage 4.0 software (Thorlabs, USA).

### Data analyses for electrophysiology and two-photon imaging

Analysis of hippocampal field potentials was conducted using Clampfit 10.3 software (Molecular Devices, USA). fEPSPs were quantified by measuring the slope of their linear part on the rising phase of the response and normalized to the average of the baseline. The collected image series were analyzed using the Fiji imaging software (http://fiji.sc)(Schindelin et al., 2012). Cell somas located in the CA1 region of *stratum* pyramidale (or neuronal extensions located in CA1 *stratum radiatum/oriens*) were defined manually and selected as regions of interest (ROIs). The fluorescence intensity trace, F, was normalized to the mean baseline intensity before stimulation, F(0), and the average normalized fluorescence intensity calculated as mean(F/F(0)). Data are represented using GraphPad Prism version 8 (GraphPad Software, La Jolla, CA).

### Hippocampal dissociated cultures

Brains isolated from 4-6 prenatal E19 Wistar rat embryos of mixed gender (Charles River, Wilmington, MA) were placed in ice-cold dissection medium [HBSS supplemented with 30 mM glucose, 10 mM HEPES (pH=7.4), 1 mM sodium pyruvate, 100 u/mL penicillin, 100 µg/mL streptomycin]. The two hemispheres were parted by a sagittal cut and cerebellum removed. The meninges were removed before carefully dissecting out the hippocampus from each forebrain using two small forceps. Isolated hippocampi were collected and kept in ice-cold dissection medium. The hippocampi were treated with papain (Worthington, Lakewood, NJ)(84.8 µgP/mL in dissection medium) for 20 min at 37°C, triturated (2×10 times) using differentially fire-polished Pasteur pipettes, and filtered through a 70 µm cell strainer. The cells were seeded onto poly-L-lysine-coated 25 mm glass coverslips emerged in Neurobasal medium (supplemented with 2% glutamax, 1% pen/strep, 4% FBS and 2% B27 supplement) (Thermo Fischer Scientific, Waltham, MA) at a density of 100,000 cells/coverslip. Cell cultures were grown at 37°C with 5% CO_2_ and 95% humidified atmosphere. After 24 hrs, the growth medium was substituted with serum-free medium, and cells were grown for 12-18 DIV.

The neurons were co-transfected with cytosolic located super-ecliptic pHluorin, cyto-pHluorin (Rathje et al., 2013), and R-GECO1.2 (Campbell R., Addgene plasmid 45494)(Wu et al., 2013) via the Lipofectamine2000 system (Thermo Fischer Scientific). Briefly, transfection was carried out in Neurobasal medium (supplemented with 2% glutamax and 1% pen/strep). Each DNA construct was added in a quantity of 1 µg/well together with Lipofectamine2000 in a ratio of 1:3 (w/v). After incubation for 3 hrs at 37°C, the transfection medium was exchanged for normal growth medium. Imaging experiments were performed 48 hrs after transfection.

### Fluorescence widefield imaging

Time-lapse images with a frame interval of 1 min were acquired with a Zeiss LSM 510 inverted laser scanning microscope, using a 63x/1.4 oil immersion objective and ZEN2009 imaging software (Zeiss, Oberkochen, Germany). Coverslips were mounted into a QE-1/RC-40LP flow chamber (Warner instruments, Hamden, CT). Excitation light was separated with a 488/543/633 nm beam-splitter, and a 488/543 nm dichroic mirror. Cyto-pHluorin was excited with the 488 nm laser line from an argon-krypton laser (Lasos, Jena, Germany), and emitted fluorescence was detected using a 505-550 nm band-pass filter (Zeiss). R-GECO1.2 was excited with the 543 nm laser line from a helium-neon laser (Lasos), and emitted light detected using a 560 nm long-pass filter. The 633-nm laser line reflection system was applied for auto-focus to avoid focus drift.

Time-lapse images with a frame interval of 2-5 s were acquired with a Nikon eclipse Ti microscope, using a 100x/1.49 oil-immersion objective and NIS-Elements software (Nikon, Tokyo, Japan). Coverslips were mounted into a RC-21BDW microscope flow chamber (Warner instruments). Cyto-pHluorin was excited with the 488 nm laser line from an LED sapphire laser (Coherent Inc., Santa Clara, USA) using a 491 nm mirror and a 475/35 nm band-pass filter, and emitted light was detected with a 525/50 nm band-pass filter (AHF Analysentechnik, Tübingen-Pfrondorf, Germany). R-GECO1.2 was excited with the 561 nm laser line from a LED sapphire laser (Coherent Inc.), and emitted light was detected with a 630/75 nm band-pass filter (AHF Analysentechnik, Tübingen-Pfrondorf, Germany).

All experiments were performed at room temperature (RT). During baseline conditions, cells were continuously perfused with conventional aCSF buffer [120 mM NaCl, 5 mM KCl, 2 mM CaCl_2_, 2 mM MgCl_2_, 25 mM HEPES (pH=7.4), 30 mM glucose, 1 µM TTX (Tocris, Bio-Techne, Minneapolis, MN)] with a flow rate of 1 mL/min using gravity flow and FR-55S control valve or VC-6 six channel perfusion valve control system (Warner instruments). L-glutamate and NMDA stimulations were performed in reduced Mg^2+^ (0.3 mM MgCl_2_) buffer. AMPA stimulations were carried out in low Mg^2+^ buffer for experiments with a frame interval of 1 min, and in conventional aCSF buffer for experiments with a frame interval of 3-5 s. KCl-mediated depolarization and all other treatments were carried out in the conventional aCSF buffer. Stimulation and depolarization protocols were performed for 3 min and were followed by perfusion with aCSF buffer until recovery of baseline fluorescence. The final concentration of the reagents were as follows: L-glutamate (10 µM), NMDA (20 µM NMDA (Tocris), 10 µM glycine), AMPA (100 µM, Tocris), DL-APV (100 µM, Tocris), NBQX (50 µM, Hello Bio, Bristol, UK), DHPG (100 µM (R,S)-3,5-DHPG, Hello Bio, Bristol, UK), verapamil (100 µM), NASPM (30 µM, Tocris), CCCP (5 µM, Tocris), GABA (50 or 200 µM, Sigma), BAPTA-AM (20 µM, Thermo Fischer Scientific), ionomycin (10 µM, Hello Bio). The effect of pyruvate was tested by substituting glucose (30 mM) for sodium pyruvate (60 mM) in baseline and stimulation buffers, or by addition of 10 mM sodium pyruvate to the conventional aCSF buffer. Dependence on extracellular Ca^2+^ was tested by excluding CaCl_2_ from baseline and stimulation buffers.

### Data analyses for widefield imaging

The collected image series were analyzed using the Fiji imaging software (http://fiji.sc)(Schindelin et al., 2012). Cell somas were defined manually and selected as region of interest (ROI). The background-corrected intensity trace, F, was then normalized to the mean baseline intensity before treatments, F(0). Data are presented using GraphPad Prism version 8. Statistical analyses of individual responses were based on the maximum fold change in fluorescence induced by stimulations, given as the logarithmic values of the normalized R-GECO1.2 and cyto-pHluorin responses, log(F/F(0)). The data were evaluated by Brown-Forsythe and Welch’s ONE-way ANOVA tests with Dunnett’s T3 multiple comparisons (>two groups), or by Student’s *t* tests (two groups). Analyses of response ratios were based on the logarithmic values of inversed cyto-pHluorin to R-GECO normalized responses, log((1/ΔCyto-pHluorin)/ΔR-GECO1.2). Response ratio data were evaluated by Kruskal-Wallis test of column with Dunn’s multiple comparisons (>two groups), or Mann-Whitney tests (two groups).

Heatmaps and cross-correlation analysis were performed and plotted with RStudio version 4.3.2, using *Heatmap* function from *ComplexHeatmap* package for heatmap representations and *ccf* function from *Stats* package and *ggplot* from *ggplot2* package for cross-correlation analysis.

### DNA constructs and expression in Xenopus oocytes

cDNAs encoding GluN1-1a (hereafter GluN1), GluN2A, and the flip isoform of GluA1 were generously provided by Dr. S. Heinemann (Salk Institute, La Jolla, CA), or Dr. S. Nakanishi (Osaka Bioscience Institute, Osaka, Japan).

For expression in *Xenopus laevis* oocytes, cDNAs were linearized and used as templates to synthesize cRNA using T7 RNA polymerase mMessage mMachine kit (Ambion, Life Technologies, Paisley, UK). Defolliculated stage V-VI oocytes ready to inject were obtained from EcoCyte Biosciences (Castrop-Rauxel, Germany). The oocytes were coinjected with cRNAs encoding GluN1 and GluN2A in a 1:2 ratio, or cRNA encoding GluA1 and maintained at 18°C in Barth’s solution containing 88 mM NaCl, 1 mM KCl, 2.4 mM NaHCO_3_, 0.82 mM MgSO_4_, 0.33 mM Ca(NO_3_)_2_, 0.91 mM CaCl_2_, 10 mM HEPES (pH 7.5 with NaOH) supplemented with 100 IU/ml penicillin, 100 μg/ml streptomycin and 100 μg/ml gentamycin (Invitrogen, Life Technologies, Paisley, UK).

### Two-Electrode Voltage-Clamp Recordings

Two-electrode voltage-clamp (TEVC) recordings were performed on *Xenopus* oocytes at room temperature 1-3 days post-injection using an OC-725C TEVC amplifier (Warner Instruments). Glass electrodes had at tip resistance of 0.5-2.5 MΩ and were pulled from thin-walled glass capillary tubes (World Precision Instruments, Hertfordshire, UK) using a PC-10 puller (Narishige, East Meadow, NY). Voltage and current electrodes were filled with 0.3 and 3 M KCl, respectively. During recordings, oocytes were placed in a recording chamber and perfused with the extracellular recording solution comprised of 90 mM NaCl, 1 mM KCl, 10 mM HEPES, 0.5 mM BaCl_2_ and 0.01 mM EDTA (pH 7.4 with NaOH) or pyruvate-buffer 60 mM Na-Pyruvate, 30 mM NaCl, 1 mM KCl, 10 mM HEPES, 0.5 mM BaCl_2_ and 0.01 mM EDTA (pH 7.4 with NaOH). Current responses were recorded at a holding potential of −40 mV or −60 mV for recordings at GluN1/N2A and GluA1 receptors, respectively. Compounds were dissolved in extracellular recording solution or pyruvate-buffer and applied to the oocyte by gravity-driven perfusion using a computer controlled 8-modular-valve positioner (Digital MVP, Hamilton, Reno, NV). Glycine, L-glutamate, *N*-methyl-D-aspartate (NMDA) and all buffer reagents were purchased from Sigma-Aldrich (St. Louis, MO).

### Data analyses for concentration-response curves

Data were analyzed using GraphPad Prism 5. Agonist concentration-response data for individual oocytes were fitted to the Hill equation, 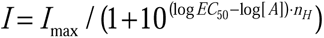. I_max_ is the maximum current in response to agonist, EC_50_ is the concentration of agonist that produces half-maximum activation, [A] is the concentration of agonist, and *n*_H_ is the Hill coefficient. The logEC_50_ from the individual oocytes were used to calculate mean and S.E.M. For graphical presentation, the data for individual oocytes were normalized to the maximum current response in the same recording and averaged. The averaged data points were then fitted to the Hill equation and plotted together with the resulting curve.

## Results

### LTP inducing theta-burst stimulation causes acidification in postsynaptic CA1 neurons

Cytosolic pH shifts in neurons have previously been reported to occur in hippocampal slices in response to chemical or electrical seizure-like stimulations (Xiong and Stringer, 2000; Xiong et al., 2000; Raimondo et al., 2012). Moreover, data suggest that pH shifts in neuronal microdomains can also occur in response to physiological activation *in vivo* electrical stimulation *ex vivo* (Zhang et al., 2010; Rossano et al., 2013).

We investigated whether LTP-inducing stimulation in hippocampal slices would lead to neuronal acidification concurrent with activity-induced cytosolic Ca^2+^ increase.To investigate this, we stereotactically injected AAVs encoding biosensors for Ca^2+^, jRGECO1a (Dana et al., 2016) and for cytosolic pH, cyto-pHluorin (Rathje et al., 2013) into the CA1 region of four-week-old mice (Fig.1A,B). After three weeks, biosensor fluorescence could be detected from both somas and extensions of CA1 pyramidal neurons in acutely isolated hippocampal slices by use of 2-photon microscopy (Fig.1A-D). LTP was induced by theta-burst stimulation (20xTBS) at CA3-CA1 synapses and confirmed by the increase of fEPSP slope in the CA1 region (Fig.1E). Concomitantly, biosensor responses to LTP-inducing stimulation were evaluated (Fig.1F-H). LTP-inducing stimulation at the Schaffer collaterals elicited a rapid and reversible (~4-5 s duration) rise in mean jRGECO1a fluorescence, indicating a rise in cytosolic Ca^2+^. Importantly, the stimulation also caused a clear decrease in mean cyto-pHluorin fluorescence in the somas of CA1 neurons (Fig.1 F-H), indicating cytosolic acidification. Although the time to peak for jRGECO1a responses was significantly shorter than for cyto-pHluorin responses (1.7-4.3 s compared to ~11-23 s), we could not distinguish a difference in their initiation time points (both ~1-2 s)(Fig.S1). The complete datasets for all slice experiments, including responses from individual cell somas, are shown in appendix 1.

**Figure 1.**
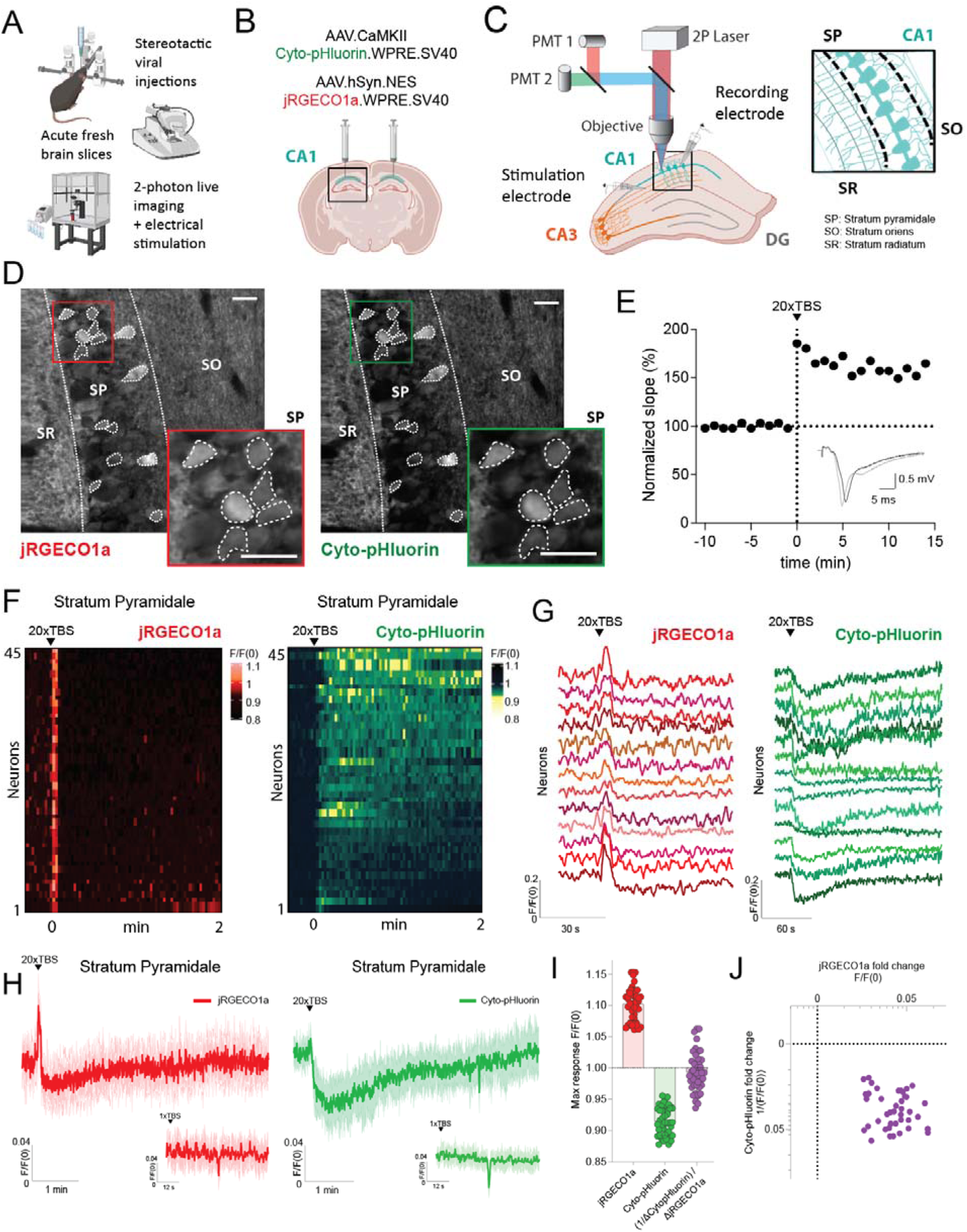
LTP-inducing theta-burst stimulation (TBS) of hippocampal Schaffer collaterals induces cytosolic acidification in postsynaptic CA1 neurons. (*A*) Graphical depiction of the experimental design (*B*) Representation of the injection sites of the dual AAV strategy for expression of the jRGECO1a Ca^2+^ sensor and cyto-pHluorin pH sensor; AAV.CaMKII.Cyto-pHluorin.WPRE.SV40 and AAV.hSyn.NES.jRGECO1a.WPRE.SV40. The viruses were injected bilateral on the CA1 region of the hippocampus. (*C*) Illustration of viral experimental setup. Acute hippocampal slices were prepared from C57bl6N mice with the AAV-mediated neuron-specific expression of the cytosolic biosensors jRGECO1a and cyto-pHluorin. Fluorescence intensities were recorded from the hippocampal CA1 region using 2-photon microscopy before, during and after LTP-inducing 20xTBS or 1xTBS stimulation to the Schaffer collaterals. Successful LTP establishment at CA3-CA1 synapses by 20xTBS stimulation was assessed by continuous recording of evoked field potentials in the CA1 region. The insert outlines the different hippocampal layers. (*D*) Representative median filtered 2-photon microscopy images of a part of the hippocampal CA1 region, showing the expression of the Ca^2+^-sensor, jRGECO1a, and the pH sensor, cyto-pHluorin. (Scale bar, 20 µM). The somatic region of CA1 neurons, defined by the dotted lines, were used in fluorescence analyses. The inserts (bottom right side for each image) show a magnification of a sample of analyzed somas. (*E*) Example trace of normalized field potential recordings before and after LTP induction through TBS stimulation. The inset displays representative fEPSP recordings before (black) and after (grey) LTP induction. (*F*) Heatmap representation of jRGECO1a and Cyto-pHluorin responses upon the 20XTBS stimulation. (*G*) Individual representative traces from neuronal responses within the same slice shown in (D) for jRGECO1a and Cyto-pHluorin. (*H*) Representative time course of the normalized mean fluorescence (mean F/F0) for jRGECO1a (red) and cyto-pHluorin (green) from the sample of CA1 somas shown in (D) stimulated with 20xTBS. The time course shows the change in fluorescence before, during and after LTP induction. Control traces from CA1 somas stimulated with 1xTBS shown at the bottom right of each of the 20xTBS representative traces. The shaded area illustrates the standard deviation from the mean response. (*I*) Individual maximal fluorescence responses induced by the LTP-inducing stimulation. The maximal response values reflect the maximal fold change in fluorescence for jRGECO1a and cyto-pHluorin in individual somas, recorded from three different slices. Ratio between both maximal responses is also plotted as (1/ΔCyto-pHluorin)/ ΔjRGECO1a. (*J*) Comparison of the maximal responses in individual experiments is shown in a dotplot. Data are shown on log10-scales. (N = 45 cells). SR: Stratum Radiatum, SP: Stratum Pyramidale, SO: Stratum Oriens.

The mean jRGECO1a fluorescence decreased to below baseline following the initial peak. The kinetics of this fluorescence decrease are consistent with reduced jRGECO1a fluorescence at low pH levels. In two out of three slices, the jRGECO1a fluorescence recovered to the original baseline level within 4-5 min after stimulation onset, while the mean cyto-pHluorin fluorescence recovered to baseline levels within 4-7 min after stimulation onset. In one slice, we saw no post-stimulatory fluorescence recovery for either biosensor within the duration of the recording (~7 min).

To assess if the LTP-inducing stimulation resulted in comparable responses in all analyzed cell somas, we plotted the maximal fluorescence responses for cyto-pHluorin and jRGECO1a for individual cells, as well as the ratio of both responses per neuron (Fig.1I-J). These data indicate homogenous activation and acidification in all CA1 neurons in the recorded area. Similar fluorescent changes were seen in the stratum *radiatum* and *oriens*, containing CA1 apical and basal dendrites, respectively (Fig.S2). This indicates a general cytosolic acidification and Ca^2+^ rise in the CA1 neurons in response to the LTP-inducing stimulation. Additionally, a 1xTBS stimulation to the Schaffer collaterals was performed as a control, which did not induce a Ca^2+^ nor pH response (Fig.1H).

In summary, these results demonstrate that hippocampal neurons undergo an acidification period upon LTP-stimulation. Moreover, we show that the dual-biosensor approach can detect cytosolic acidification, together with the Ca^2+^ rise, in hippocampal neurons in an intact synaptic network from LTP-inducing stimulation and not only by epileptic activity as previously shown (Raimondo et al., 2012).

### Spontaneous neuronal activity in dissociated hippocampal cultures induces cytosolic acidification

Next, we investigated whether spontaneous neuronal activity was sufficient to trigger acidification in the absence of any external electrical or pharmacological manipulations. To explore this, we co-transduced dissociated hippocampal cells with both a Ca^2+^ sensor (jRGECO1a) and a pH sensor (cyto-pHluorin) (Fig.2A-B). Numerous studies have demonstrated the occurrence of spontaneous neuronal activity in hippocampal cultures (Charlesworth et al., 2015, Cohen et al., 2008, Ivenshitz et al., 2010, Pimashkin et al., 2011, Sombati et al., 1995). We used a widefield microscope to detect fluorescence changes in dissociated neurons, focusing on the somatic region for analysis. We noted a range of spontaneous activity in cultures, from low-amplitude, frequency, and duration Ca^2+^ transients to robust synchronous network activity with high frequency and sustained Ca^2+^ increases (Fig.1C). Notably, changes in pH over time showed similar patterns of fluctuations to the ones observed for Ca^2+^ (Fig.2C,D,F). Additionally, both Ca^2+^ and pH oscillations were blocked by bath application of tetrodotoxin (TTX), indicating that both signals are activity-dependent (Fig.2E).

**Figure 2.**
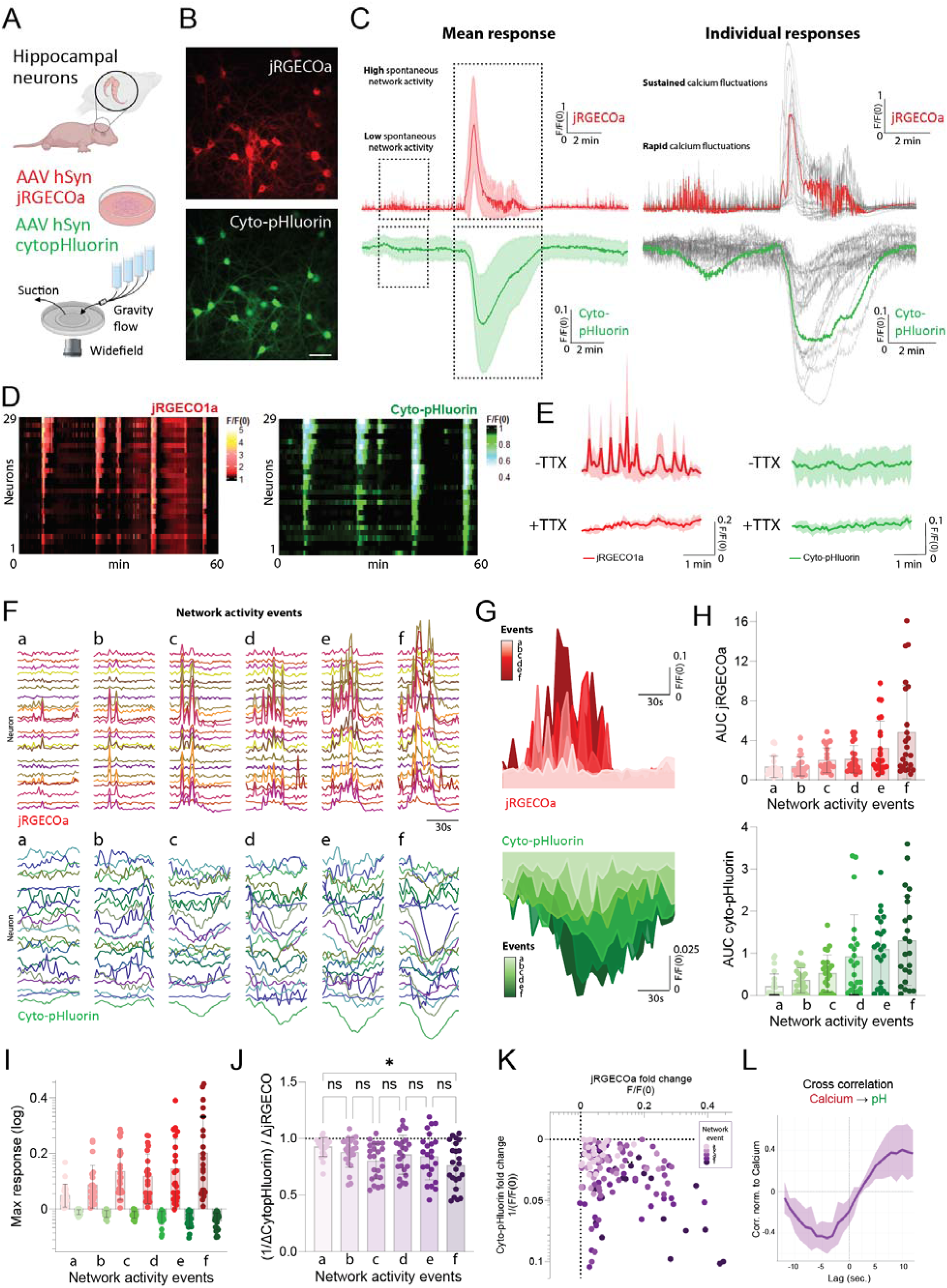
Spontaneous neuronal activity in dissociated hippocampal cultures induces cytosolic acidification. (*A*) Graphical depiction of the experimental design. (*B*) Representative median filtered widefield microscopy images of hippocampal cultured neurons, showing the expression of the Ca^2+^-sensor, jRGECO1a, and the pH sensor, cyto-pHluorin. (Scale bar, 40 µM). (*C*) Left, normalized mean fluorescence (mean F/F0) for jRGECO1a and Cyto-pHluorin showing spontaneous activity events of a neuronal culture. Dotted-line boxes indicate two different types of activity. Shaded area represents ± SD. Right, individual neuronal traces from the same culture, shown in grey. Highlighted Ca^2+^ (red) and pH (green) traces correspond to an individual neuron serving as an example. (N = 29 cells) (*D*) Heatmap representation of jRGECO1a and Cyto-pHluorin signals over time. (N = 29 cells) (*E*) Representative time course of the normalized mean fluorescence (mean F/F0) for jRGECO1a (red) and cyto-pHluorin (green) in the presence and absence of tetrodotoxin (TTX). Shaded area indicates ±SD. (N = 33 cells) (*F*) Individual normalized neuronal traces for jRGECO1a and Cyto-pHluorin from the same neuronal population at different selected time points (a,b,c,d,e and f) where network activity events spontaneously occurred, ordered by intensity where “a” corresponds to the weakest activity event and “f” to the strongest. Same period length around each event was selected for further analysis. (N = 25 cells). (*G*) Mean traces with colorized area under the curve (AUC) corresponding to each network activity event shown in (F) from the same neurons at different time points for jRGECO1a and Cyto-pHluorin. (*I*) Individual AUC measurements plotted from each neuron at the different network activity events (a-f) for jRGECO1a and Cyto-pHluorin. (*H*) The maximal response for jRGECO1a and Cyto-pHluorin plotted for each event. Values reflect the maximal fold change in fluorescence for jRGECO1a and cyto-pHluorin in individual somas (J) Ratio from maximal responses shown in (I) for each event plotted as (1/ΔCyto-pHluorin)/ ΔjRGECO1a. Experiments are evaluated by Kruskal-Wallis test of column with Dunn’s multiple comparisons test *P≤0.05 (*K*) Comparison of the maximal responses for each neuron for each event shown as a dotplot. Data are shown on log10-scales. (*L*) Cross-correlation analysis between Ca^2+^ and pH fluctuations for all neurons in all the events normalized to the Ca^2+^ events. (N = 25 cells).

To further explore the relationship between activity intensity and acidification, we isolated individual network activity events within the same neuronal population occurring over time and ordered them based on their Ca^2+^ rises as a proxy for the neuronal activity intensity (Fig.2F). When comparing the Ca^2+^ signal per event to its pH response, both quantified as area under the curve, we found that pH exhibited a concentration-response-like pattern to Ca^2+^, with increased Ca^2+^ influx giving rise to more pronounced intracellular acidification (Fig.2G-H). This correlation remained constant across events when analyzing the ratios of maximal Ca^2+^ and pH responses per neuron, showing a significant difference only between the ratios of the strongest and weakest events (Fig.2I-K). Cross-correlation analysis further supported this relationship, indicating a lag of approximately ~8 seconds between the maximal amplitude Ca^2+^ event and subsequent maximal pH response (Fig.2L). From these experiments we could not determine whether this lag was due to a lag in the onset of the pH response or due to slower kinetics.

In conclusion, our findings demonstrate the presence of activity-induced cytosolic acidification, not only during robust stimulation paradigms, but also during spontaneous events, highlighting a concentration-response relationship between cytosolic Ca^2+^ and H^+^ increase.

### Glutamate receptor activation causes more robust cytosolic acidification than simple depolarization

Next, we aimed to dissect what component of the activity is causing this phenomenon. Administration of glutamate to dissociated hippocampal cultures was previously shown to induce intracellular acidification in neurons (Hartley and Dubinsky, 1993; Wang et al., 1994). We co-transfected dissociated hippocampal cells with a Ca^2+^ sensor, R-GECO1.2 (Wu et al., 2013), and the pH sensor, cyto-pHluorin (Rathje et al., 2013) and first assessed the neuronal responses induced by bath application of 20 µM glutamate for 3 min (Fig.3A-B). In line with previous reports, glutamate induced an immediate and robust reduction in cyto-pHluorin fluorescence, indicating cytosolic acidification, which recovered to baseline levels within 20-40 min (Fig.3C). The acidification was accompanied by a rapid rise in R-GECO1.2 fluorescence, indicating an increase in cytosolic Ca^2+^, lasting for 3-12 min before typically shifting to below baseline and eventually reaching a new, slightly higher steady-state after 15-25 min. The transient shift to below baseline fluorescence may, at least partly, be attributed to the pH sensitivity of R-GECO1.2, which shows reduced fluorescence below pH 7 (Wu et al., 2013)(Fig.S3). Under physiological conditions, neuronal pH homeostasis is dependent on the bicarbonate buffer system (reviewed in (Chesler, 2003)). Indeed, experiments confirmed a reduced acidification relative to Ca^2+^ rise in neurons placed in a bicarbonate buffered system compared to a HEPES buffered system (Fig.S4). Nonetheless, significant acidification was evident in both buffer systems.

**Figure 3.**
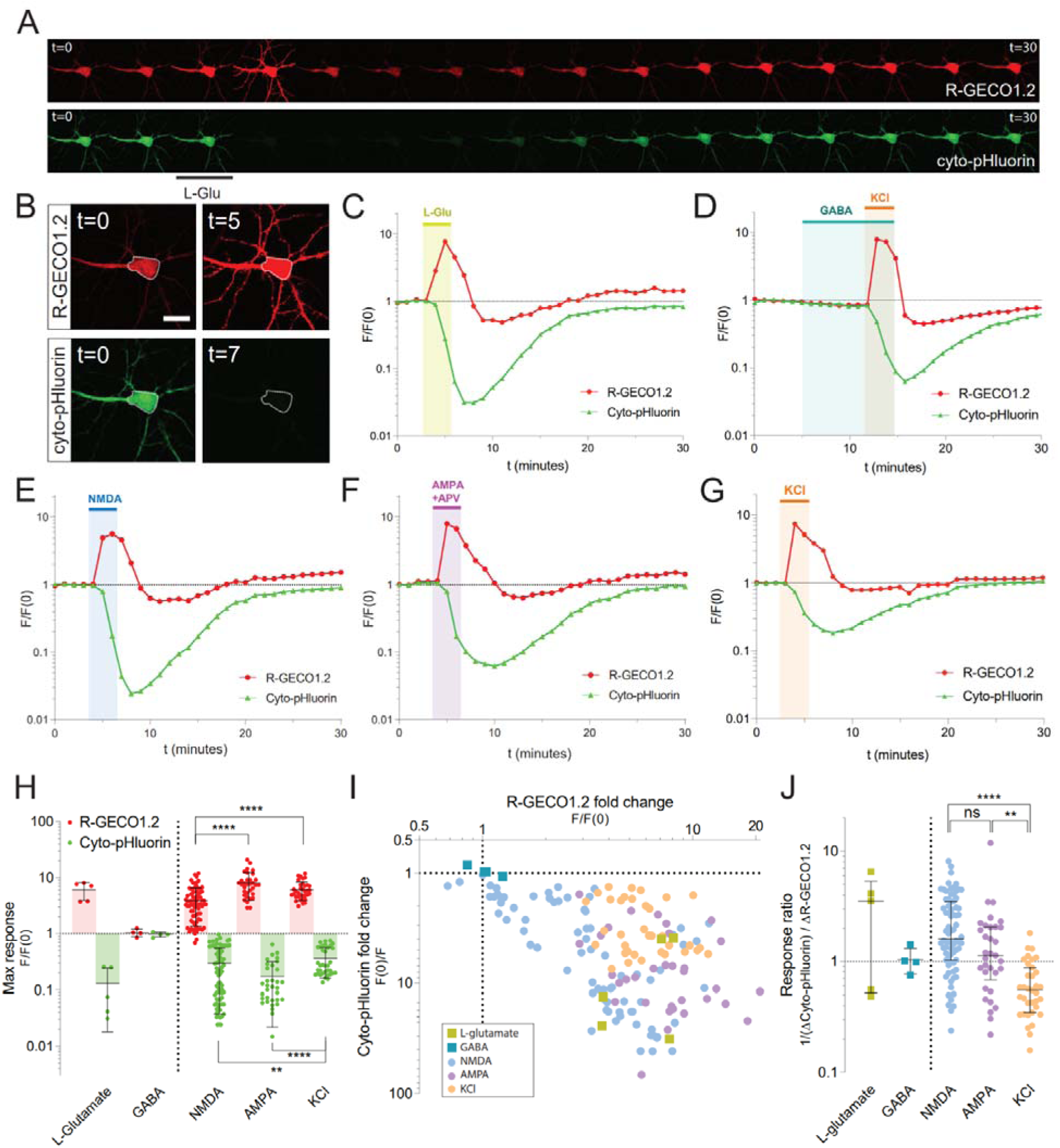
Characterization of the cytosolic acidification in dissociated hippocampal neurons induced by chemical stimulation or depolarization. *(A)* Representative image sequence (2 minutes interval for 30 minutes) of a hippocampal neuron co-expressing R-GECO1.2 (red) and cyto-pHluorin (green). (Scale bar, 20 μm). Images show the change in fluorescence in response to L-glutamate. *(B)* Representative magnified images from (A) before and after L-glutamate stimulation where dotted line shows the area measured. *(C-G)* Representative time course of the normalized fluorescence (F/F0) from the somatic region (see FigS1A), in response to chemical stimulation or depolarization, plotted on a log10-scale. L-glutamate (10 μM), GABA (50 µM for 10-15 min, or 200 µM for 10-20 min), NMDA (20 μM NMDA, 10 μM glycine), AMPA (100 μM AMPA, 100 μM APV) or high KCl (60 mM) was applied at the indicated time points for 3 min. (H) Comparison of the maximal responses induced by the stimulation protocols. The maximal response values reflect the maximal fold change in fluorescence for R-GECO1.2 and cyto-pHluorin in individual experiments. Data are shown on a log10-scale with mean ± SD. Log-transformed R-GECO1.2 or cyto-pHluorin responses from NMDA, AMPA and KCl experiments (log(F/F_0_)) are evaluated by Brown-Forsythe and Welch ONE-way ANOVA tests, with Dunnett’s T3 multiple comparisons test, *P≤0.1, ****P≤0.0001. AMPA data include AMPA responses in presence and absence of APV. KCl data include KCl responses in presence of APV and NBQX. *(I)* Comparison of cyto-pHluorin to R-GECO1.2 maximal response ratios for individual experiments. Ratios are calculated with inversed values of cyto-pHluorin responses (F(0)/F), and are shown on a log10-scale with medians and interquartile ranges. Data from NMDA, AMPA and KCl experiments are evaluated by Kruskal-Wallis test of column with Dunn’s multiple comparisons test, *P≤0.1, ****P≤0.0001. L-glutamate (N = 5 cells), GABA data (N = 4 cells), NMDA (N = 76 cells), AMPA (N = 33 cells), KCl (N = 28 cells). *(J)* x-y plot showing the maximal cyto-pHluorin response as a function of the maximal R-GECO1.2 response for each tested cell. The maximal response values reflect the maximal fold change in fluorescence for R-GECO1.2 and cyto-pHluorin in individual experiments, with cyto-pHluorin responses plotted as inversed values (F(0)/F). Data is a summary of the initial stimulations obtained throughout the rest of study and are shown on log10-scales.

Administration of GABA to dissociated hippocampal cultures has previously been shown to induce neuronal acidification (Pasternack et al., 1993). To rule out that activity-dependent GABA release could result in GABA_A_ receptor-mediated cytosolic acidification in our setup and influence our results, we tested the effect of bath administrating 50 µM or 200 µM GABA for 10-20 min (Fig.3D and Fig.S5). Our results showed that GABA had no apparent effect on cyto-pHluorin or R-GECO1.2 fluorescence, while a control stimulation with 60 mM KCl induced clear responses from both biosensors. Previous GABA-induced acidification studies have been performed in a bicarbonate-buffered system, likely explaining this discrepancy (Pasternack et al., 1993).

Together, these results indicated that a HEPES buffered imaging setup would allow us to confine our study to cytosolic pH responses induced by excitatory activity and without influence of bicarbonate transport.

We next assessed the relationship between pH and Ca^2+^ in dissociated hippocampal neurons during the administration of specific glutamate receptor agonists, NMDA and AMPA, as well as during KCl-induced membrane depolarization. Each treatment was applied for three minutes. In agreement with previous reports (Irwin et al., 1994; Cheng et al., 2008; Rathje et al., 2013) NMDA (20 µM; EC_20_-chemical LTD protocol)(Erreger et al., 2007), AMPA (100 µM; saturating concentration)(Zhang et al., 2006) and depolarizing KCl buffer (60 mM KCl) all induced an immediate reduction in cytosolic pH accompanied by a rise in Ca^2+^, as indicated by cyto-pHluorin and R-GECO1.2 fluorescence responses (Fig.3E-G and Fig.S5). Stimulation of group I metabotropic glutamate receptors (mGluRs) (100 µM DHPG) did not induce cytosolic acidification (Fig.S6), indicating that the phenomenon is specific to ionotropic glutamate receptors. In a previous study investigating KCl-induced pH shifts, the onset of the cytosolic Ca^2+^ response was determined to precede the onset of acidification (5.4 sec vs. 17.4 sec) (Cheng et al., 2008). However, although the R-GECO1.2 fluorescence rise phase was steeper than for cyto-pHluorin, we were not able to discern a difference in response onsets in our setup, even at an image acquisition rate of 1 image/2-3 sec (Fig.S7A,B,D,F).

To quantitatively compare the responses induced by the different stimulation types, the maximum fold change in fluorescence for each biosensor, defined as the maximal responses, were grouped according to stimulation protocol (Fig.3H). For visual comparison, maximal responses from glutamate and GABA stimulation were also included. Evidently, the data show substantial cell-cell variability in response sizes within the groups. This variation was not abolished by increasing the temporal resolution of image acquisition to 1 image/3-5 sec (Fig.S7C,E,G), indicating that it was not caused by limitations in the acquisition rate. Statistical comparison between NMDA, AMPA and KCl mean responses, showed that NMDA and AMPA cyto-pHluorin responses were larger than for KCl, whereas NMDA R-GECO1.2 responses were smaller than for AMPA and KCl. These data suggest that activation of glutamate receptors typically induces more prominent cytosolic acidification compared to membrane depolarization alone, but NMDA receptors typically induce a smaller cytosolic Ca^2+^ rise compared to AMPA receptors and membrane depolarization^1^. Correlation analyses showed a good overall correlation between cyto-pHluorin and R-GECO1.2 responses (Pearson correlation coefficient, All data: P = 0.0018) (Fig.3I). The overall correlation was dominated by the NMDA data, which contributed most of the data points (Pearson correlation coefficients; L-Glu: P = 0.7273, 5 cells; NMDA: P = <0.0001, 76 cells; AMPA: P = 0.6094, 33 cells; KCl: P = 0.2766, 28 cells). Moreover, our data showed that NMDA receptors were more efficient in mediating acidification relative to the rise in Ca^2+^ than were AMPA receptors, which in turn were more efficient in mediating acidification than simple depolarization (ratios of cyto-pHluorin to R-GECO1.2 maximal responses were significantly different in the order: NMDA > AMPA > KCl) (Fig.3J). No apparent difference in pH recovery time was detected for the different treatments (Fig.S8).

### Pyruvate substitution for extracellular glucose reduces NMDA receptor efficacy

In two previous studies, NMDA- and KCl-induced cytosolic acidification was reported to arise mainly from increased neuronal glycolysis (Zhan et al., 1997, 1998). This conclusion was based on experiments that aimed to bypass glycolytic activity by substituting buffer glucose with pyruvate to directly promote mitochondrial respiration. To probe the relation between neuronal pH and Ca^2+^ under these conditions, we replicated the previous setup using dissociated hippocampal neurons co-expressing cyto-pHluorin and R-GECO1.2. Neurons were stimulated with glutamate (20 µM) for 3 min before and after an energy-equivalent substitution of pyruvate (60 mM) for glucose (30 mM) in the extracellular medium (Fig.4A,B). In agreement with the previous studies, the maximal cyto-pHluorin responses were significantly reduced during pyruvate substitution, however, our data also revealed a reduction in the maximal R-GECO1.2 responses (Fig.4C), suggesting a dependence on cytosolic Ca^2+^. To further investigate the mechanism behind the effect of pyruvate, neurons were stimulated with specific ionotropic glutamate receptor agonists, NMDA or AMPA, before and after pyruvate substitution (Fig.4D-E). Unexpectedly, pyruvate substitution completely abolished both cyto-pHluorin and R-GECO1.2 responses induced by NMDA, while it did not prevent AMPA-induced responses (Fig.4F). In contrast, repetitive NMDA administration in absence of pyruvate showed only a slight reduction in the R-GECO1.2 maximal response to the second stimulation (Fig.S9). These results suggest a role for pyruvate in direct inhibition of NMDA receptor function, rather than in preventing glycolysis-mediated acidification. In support of this notion, post-stimulatory pyruvate substitution had no apparent effect on the fluorescence recovery (Fig.S10).

**Figure 4.**
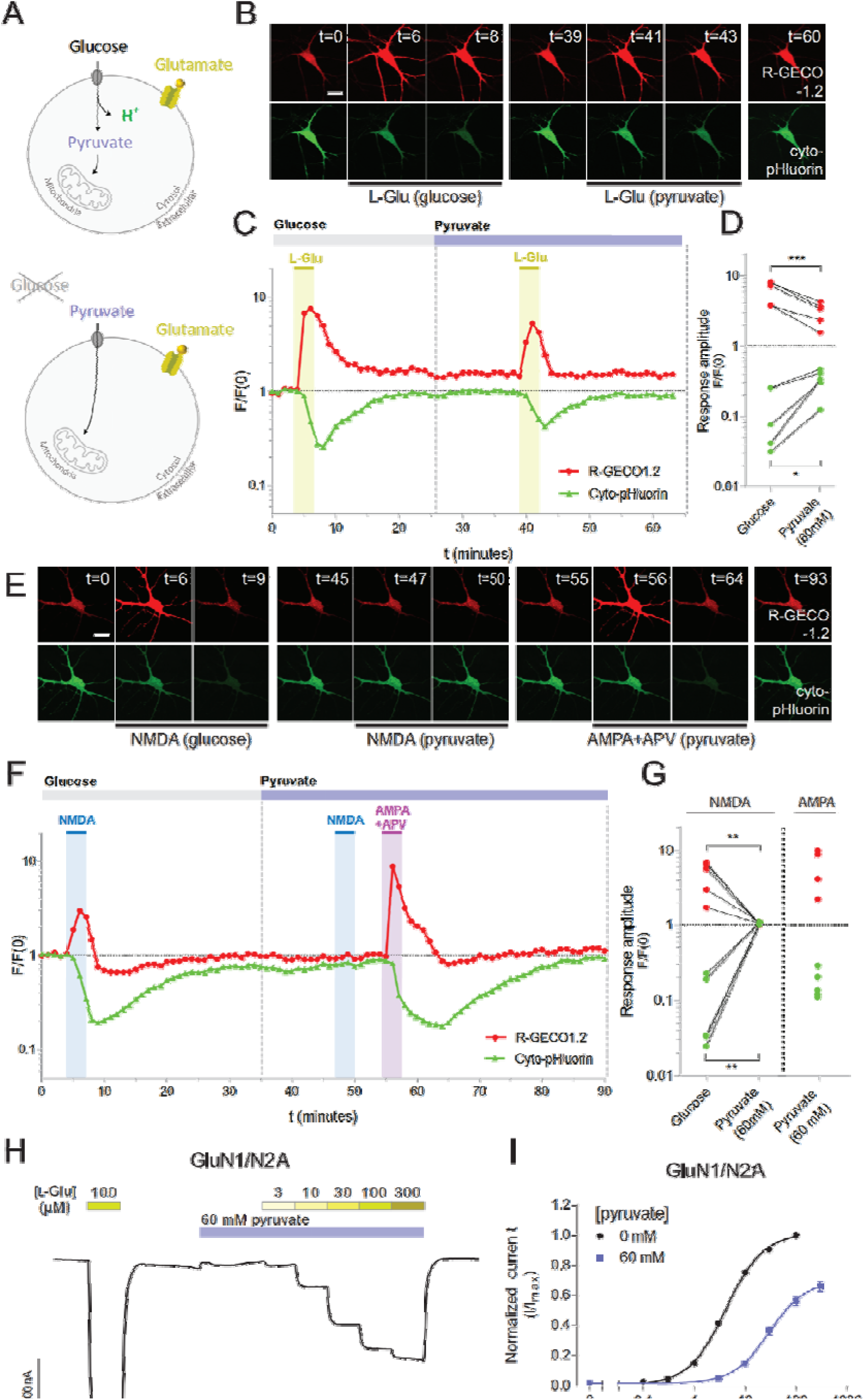
Pyruvate substitution for extracellular glucose reduces NMDA receptor conductance. (*A*) Schematic representation of expected bypass of glycolysis-produced protons by replacement of extracellular glucose for pyruvate. (*B*) Representative image sequence of a hippocampal neuron co-expressing R-GECO1.2 (red) and cyto-pHluorin (green). (Scale bar, 20 μm). Images are plotted for the indicated time points and show the change in fluorescence in response to L-glutamate (L-Glu) applied before and after pyruvate substitution (60 mM) for glucose (30 mM) in the extracellular medium. (*C*) Representative time course of the normalized fluorescence (F/F0) in the somatic region of the neuron shown in (A), plotted on a log10-scale. L-glutamate (10 μM) was applied at the indicated time points for 3 min. (*D*) Individual maximal responses induced by L-glutamate before and after pyruvate-substitution, shown on a log10-scale. Data are evaluated by ratio paired *t* test, *P≤0.05, ***P≤0.001. N = 5 cells. (*E*) Representative image sequence of a hippocampal neuron co-expressing R-GECO1.2 (red) and cyto-pHluorin (green), showing the change in fluorescence in response to NMDA or AMPA applied before and after pyruvate substitution for glucose. (F) Representative time course of the normalized fluorescence (F/F0) in the somatic region of the neuron shown in (D), plotted on a log10-scale. NMDA (20 μM NMDA, 10 µM glycine) or AMPA (100 µM AMPA, 100 µM APV) was applied at the indicated time points for 3 min. (*G*) Individual maximal responses induced by NMDA before and after pyruvate-substitution, or by AMPA after pyruvate substitution, shown on a log10-scale. Data are evaluated by ratio paired *t* test, **P≤0.01. N = 5 cells. (*H*) Representative two-electrode voltage clamp recording of current responses from recombinant GluN1/N2A receptors expressed in *Xenopus* oocytes, showing concentration-response relationship for L-glutamate in the presence of 60 mM pyruvate. All L-glutamate concentrations were co-applied with 30 µM glycine. (*I*) L-glutamate concentration-response curves at recombinant GluN1/N2A receptors, co-activated by 30 µM glycine, in the absence (0 mM) and presence (60 mM) of pyruvate. Data from 6-10 oocytes. L-glutamate: EC_50_ (pEC_50_ ± SEM) = 4.2 (5.4±0.02), R_max_ ± SEM^a^ = 100 %. L-glutamate+pyruvate (60 mM): EC_50_ (pEC_50_ ± SEM) = 32 (4.5±0.05), R_max_ ± SEM^a^ = 69±3.4 %. ^a^ The maximal responses are normalized to the maximal response of L-glutamate (100 µM at GluN1/N2) in the same recording (set to 100%).

To test if lower concentrations of pyruvate, with putative therapeutic potential for e.g. stroke treatment (Wang et al., 2009; Ryou et al., 2012), can also affect NMDA-induced responses, hippocampal neurons co-expressing cyto-pHluorin and R-GECO1.2 were stimulated with NMDA (20 µM) before and after addition of 10 mM pyruvate to the conventional aCSF buffer (Fig.S11). The lower concentration of pyruvate reduced both cyto-pHluorin and R-GECO1.2 responses, suggesting that even 10 mM pyruvate may affect NMDA receptor activity.

In order to evaluate a possible direct interaction of pyruvate with the NMDA receptor, we recombinantly expressed the GluN1/N2A NMDA receptor subtype in *Xenopus* oocytes. Using this system, the effect of pyruvate was characterized by the generation of concentration-response curves for glutamate in the absence and presence of 60 mM pyruvate (Fig.4G-H). The addition of pyruvate caused a reduction in the glutamate-induced inward current, in agreement with our results in dissociated hippocampal neurons. The recombinant NMDA receptor showed a reduction in maximal response to 69±3.4 % in the presence of pyruvate, as well as a right-shift of the concentration-response curve with an 8-fold increase in the EC_50_-value from 4.2 µM (pEC_50_ ± SEM: 5.4±0.02) compared to 32 µM (pEC_50_ ± SEM: 4.5±0.05). These results support a direct effect of pyruvate at NMDA receptors and suggest a non-competitive inhibition mechanism.

### The absence of extracellular Ca ^2+^ blunts cytosolic acidification induced by NMDA and AMPA and eliminates KCl-induced acidification

Several studies have shown that the removal of extracellular Ca^2+^ prevents stimulation-induced extracellular alkalization and cytosolic acidification in neurons (Irwin et al., 1994; Paalasmaa et al., 1994; Cheng et al., 2008; Makani and Chesler, 2010). To investigate this Ca²⁺-dependent acidification mechanism, we assessed the relation between cytosolic pH and Ca^2+^ in our system during the administration of NMDA, AMPA or KCl in the absence of extracellular Ca^2+^ (Fig.5). The absence of extracellular Ca^2+^ did not by itself affect baseline fluorescence (Fig.5B, E, H), indicating low Ca^2+^ permeability of the plasma membrane in the absence of stimuli. As expected, all stimulation-induced increases in R-GECO1.2 fluorescence were near or completely absent in the Ca²⁺-free buffer. NMDA and AMPA stimulations resulted in slight decreases in R-GECO1.2 fluorescence, which may be due to pH-dependent quenching of R-GECO1.2 fluorescence. In agreement with previous reports, stimulation of neurons in the Ca²⁺-free buffer drastically blunted the maximal cyto-pHluorin responses compared to stimulations at normal Ca^2+^ levels (Fig.5C,F,I). Interestingly, NMDA and AMPA stimulations still resulted in a slight decrease in cyto-pHluorin fluorescence, whereas no change in fluorescence was observed from KCl-induced depolarizations. This suggests that cytosolic acidification induced by ionotropic glutamate receptors relies on both Ca²⁺-dependent and independent processes, whereas depolarization-induced acidification is solely Ca²⁺-dependent.

**Figure 5.**
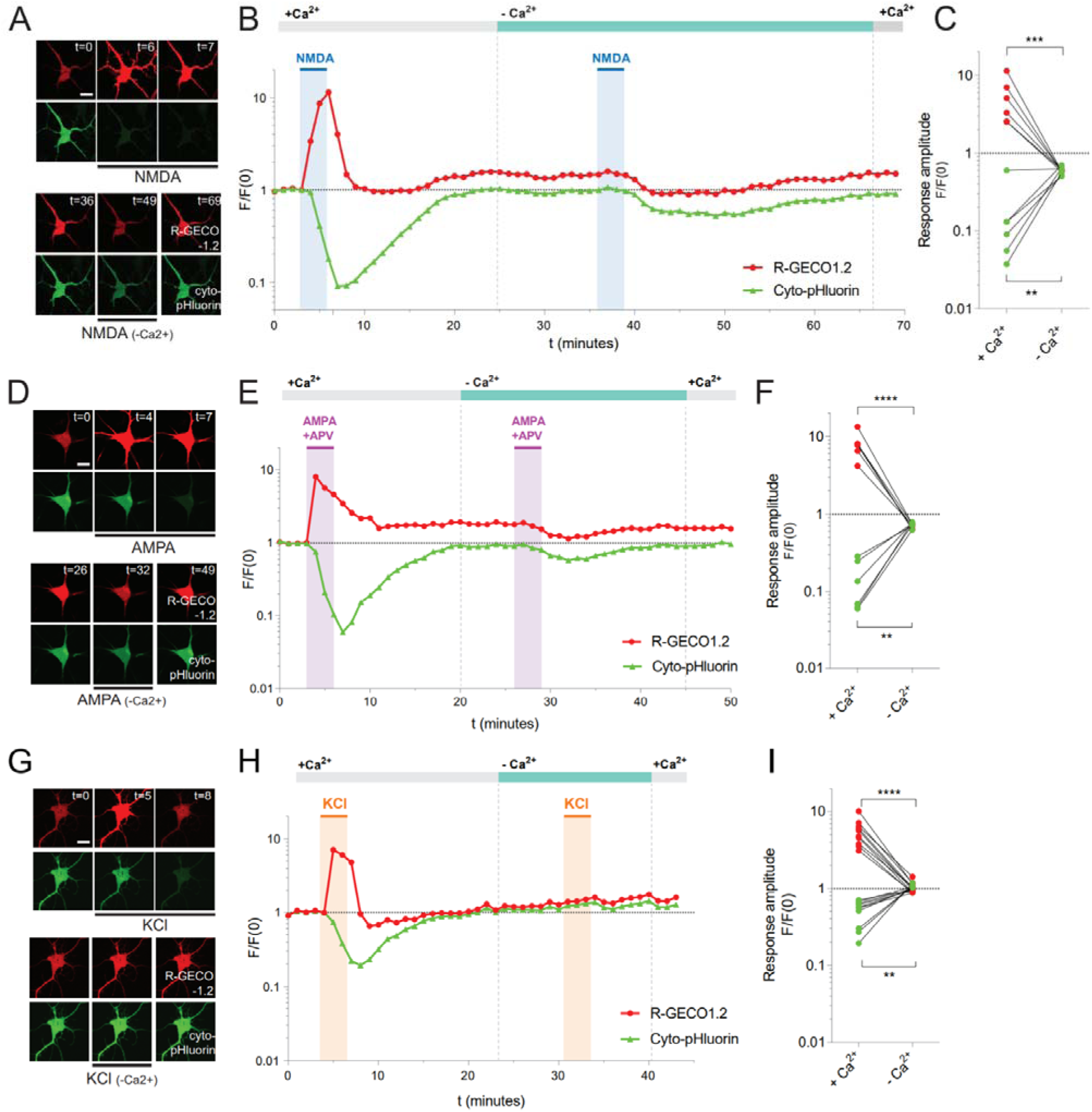
Removal of extracellular Ca^2+^ partially blunts glutamate receptor-induced cytosolic acidification, and eliminates KCl-induced cytosolic acidification. *(A, D, G)* Representative image sequences of hippocampal neurons co-expressing R-GECO1.2 (red) and cyto-pHluorin (green). (Scale bar, 20 μm). Images are plotted for the indicated time points and show the change in cell fluorescence in response to NMDA, AMPA or KCl in presence or absence of extracellular Ca^2+^. *(B, E, H)* Representative time courses of the normalized fluorescence (F/F0) in the somatic region of the neurons shown in (A, D, G), plotted on a log10-scale. NMDA (20 µM NMDA, 10 µM glycine), AMPA (100 µM AMPA, 100 µM APV), or KCl (60 mM) was applied at the indicated time points for 3 min before and after Ca^2+^ exclusion from the extracellular medium. *(C, F, I)* Individual maximal responses to the stimulation protocols in presence or absence of extracellular Ca^2+^. Data are shown on log10-scales, and are evaluated by ratio paired *t* tests, *P≤0.05, **P≤0.01, ***P≤0.001 ****P≤0.0001. NMDA (N = 5 cells), AMPA (N = 5 cells), 60 mM KCl (N = 3 cells).

### Ca^2+^ influx per se causes only weak neuronal acidification

To address if stimulation-induced cytosolic acidification could arise as a direct result of Ca^2+^ signalling or Ca^2+^ exchange for protein-bound or extracellular H^+^ (Hartley and Dubinsky, 1993; Cheng et al., 2008) we manipulated a rise in cytosolic Ca^2+^ independently of neuronal activation (Fig.6). The Ca^2+^ ionophore, ionomycin (10 µM) (Fig.6A), was applied for 1-3 min to neurons co-expressing cyto-pHluorin and R-GECO1.2 (Fig.6B-C). Ionomycin treatment induced an increase in R-GECO1.2 fluorescence, indicating cytosolic Ca^2+^ rise. In most of the tested cells, the rise in R-GECO1.2 fluorescence was persistent after ionomycin wash-out. Concomitantly with the rising phase of the R-GECO1.2 response, cyto-pHluorin fluorescence decreased slightly, indicating cytosolic acidification. The cyto-pHluorin fluorescence recovered to baseline levels within 10 minutes, irrespective of R-GECO1.2 recovery. An ionomycin treatment for 2-3 min typically induced a larger R-GECO1.2 response compared to treatments of 1 min duration, while the maximal change in cyto-pHluorin fluorescence remained low irrespective of treatment duration (Fig.S12). The maximal fluorescence responses from neurons treated with ionomycin for 2-3 min were compared to the previously obtained responses from NMDA, AMPA and KCl (Fig.6D-F). The statistical comparison showed that maximal ionomycin-induced R-GECO1.2 responses, reflecting cytosolic Ca^2+^ shifts, were comparable to R-GECO1.2 responses induced by AMPA and KCl, but not NMDA. In contrast, maximal ionomycin-induced cyto-pHluorin responses, reflecting cytosolic pH shifts, were significantly smaller than responses induced by excitatory stimulation (Fig.6D). Additional analyses, comparing biosensor response ratios across treatments, suggested that cytosolic Ca^2+^ rises induced by NMDA or AMPA are associated with larger pH drops compared to ionomycin-induced Ca^2+^ rises, whereas KCl-induced response ratios were not significantly different. These data demonstrate that neuronal Ca^2+^ influx *per se* is not sufficient to induce the substantial cytosolic acidification associated with glutamate receptor activation.

**Figure 6.**
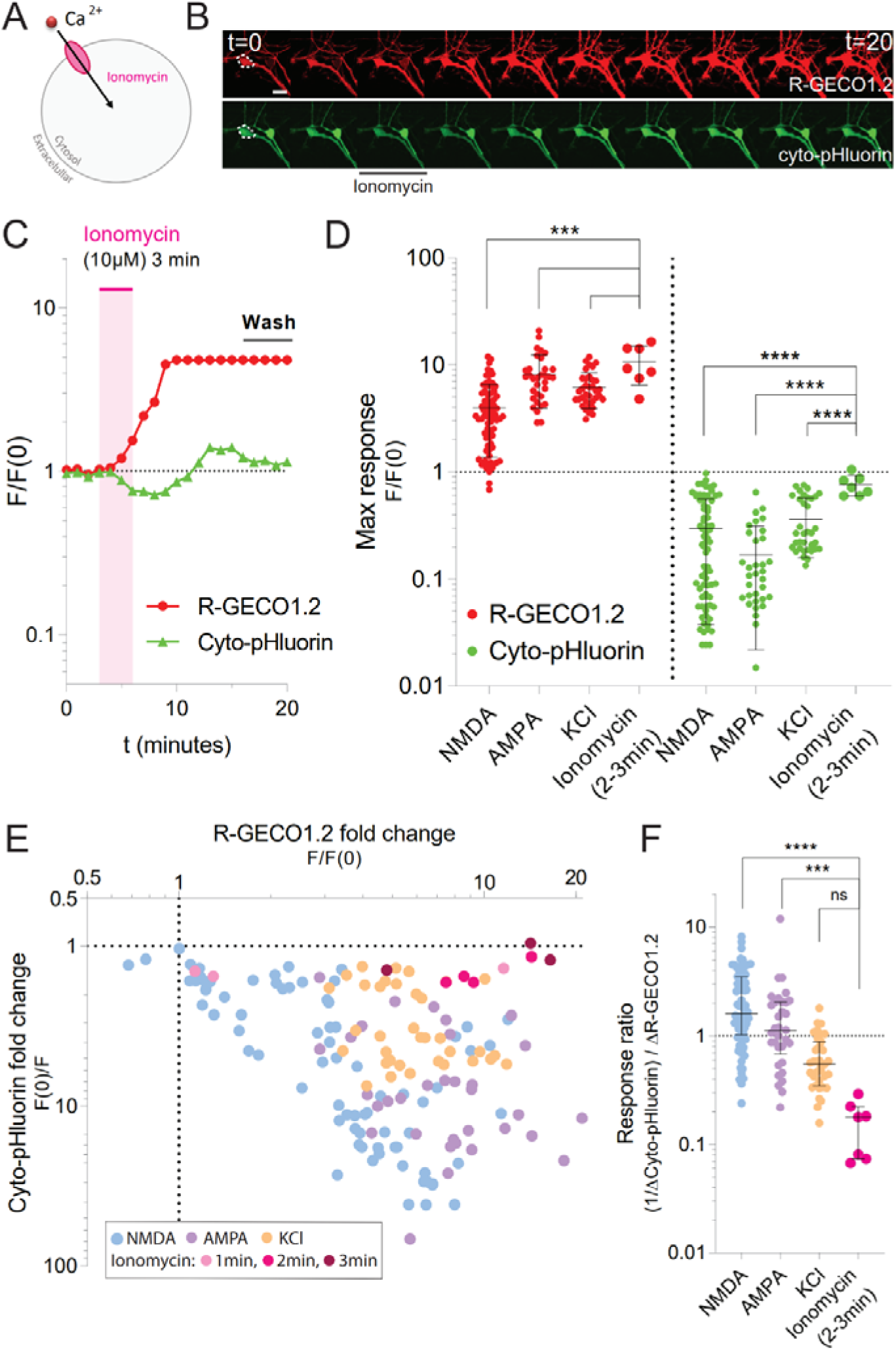
Ca^2+^ influx per se causes only weak neuronal acidification compared to the stimulation-induced acidification. (*A*) Schematic representation of the expected mechanism of action of Ionomacyn incubation. (*B*) Representative image sequence of a hippocampal neuron co-expressing R-GECO1.2 (red) and cyto-pHluorin (green). (Scale bar, 20 μm). Images are plotted for every two minutes of recording from 0 to 20 min and show the change in fluorescence in response to bath administration of the Ca^2+^ ionophore ionomycin. (*C*) Representative time course of the normalized fluorescence (F/F0) for the neuron shown in (B), plotted on a log10-scale. Ionomycin (10 µM) was applied at the indicated time point for 3 min. (*D*) Comparison of the maximal responses induced by ionomycin treatment (for 2-3 min) to the maximal responses induced by NMDA, AMPA and KCl. The maximal response values reflect the maximal fold change in fluorescence for R-GECO1.2 and cyto-pHluorin in individual experiments. Data are shown on a log10-scale with mean ± SD. Log-transformed R-GECO1.2 or cyto-pHluorin responses are evaluated by Brown-Forsythe and Welch ONE-way ANOVA tests, with Dunnett’s T3 multiple comparisons test, ****P≤0.0001. (*E*) x-y plot showing the maximal cyto-pHluorin response as a function of the maximal R-GECO1.2 response for individual cells treated with ionomycin for 1-3 min compared to stimulation with NMDA, AMPA and KCl. The maximal response values reflect the maximal fold change in fluorescence for R-GECO1.2 and cyto-pHluorin in individual experiments, with cyto-pHluorin responses plotted as inversed values (F(0)/F). Data are shown on log10-scales. (*F*) Comparison of cyto-pHluorin to R-GECO1.2 maximal response ratios for individual experiments shown in (E). Ratios are calculated with inversed values of cyto-pHluorin responses (F(0)/F), and are shown on a log10-scale with medians and interquartile ranges. Data are evaluated by Kruskal-Wallis test of column with Dunn’s multiple comparisons test, ****P≤0.0001. Ionomycin data are from 10 cells (N = 4-6 experiments for each treatment duration).

### AMPAR mediated acidification does not rely exclusively on Ca²˖-influx

As our previous results suggested that exclusion of extracellular Ca^2+^ did not fully prevent glutamate receptor-mediated acidification, we next sought to isolate a putative Ca²⁺-independent component of the receptor-dependent acid generation. We focused the following experiments on the AMPA receptors, which, in contrast to NMDA receptors, display low Ca²⁺-permeability during basal neurotransmission (Wenthold et al., 1996). The low basal expression of Ca²⁺-permeable AMPA receptors in our hippocampal culture was confirmed by the administration of an inhibitor of Ca²⁺-permeable AMPA receptors, NASPM (30 µM), which had no apparent effect on responses (neither cyto-pHluorin nor R-GECO1.2) induced by bath application of AMPA (Fig.S13). Hence, in our setup, both AMPA- and KCl-induced Ca^2+^ rises should mainly rely on the activation of voltage-gated Ca^2+^ channels.

Accordingly, AMPA-induced maximal R-GECO1.2 responses were considerably reduced by preincubation with an inhibitor of L-type voltage-gated Ca^2+^ channels, verapamil (100 µM, 30 min) (Fig.7A-D). In addition, verapamil treatment reduced but did not eliminate the AMPA-induced maximal cyto-pHluorin responses, leaving a robust acidification.

**Figure 7.**
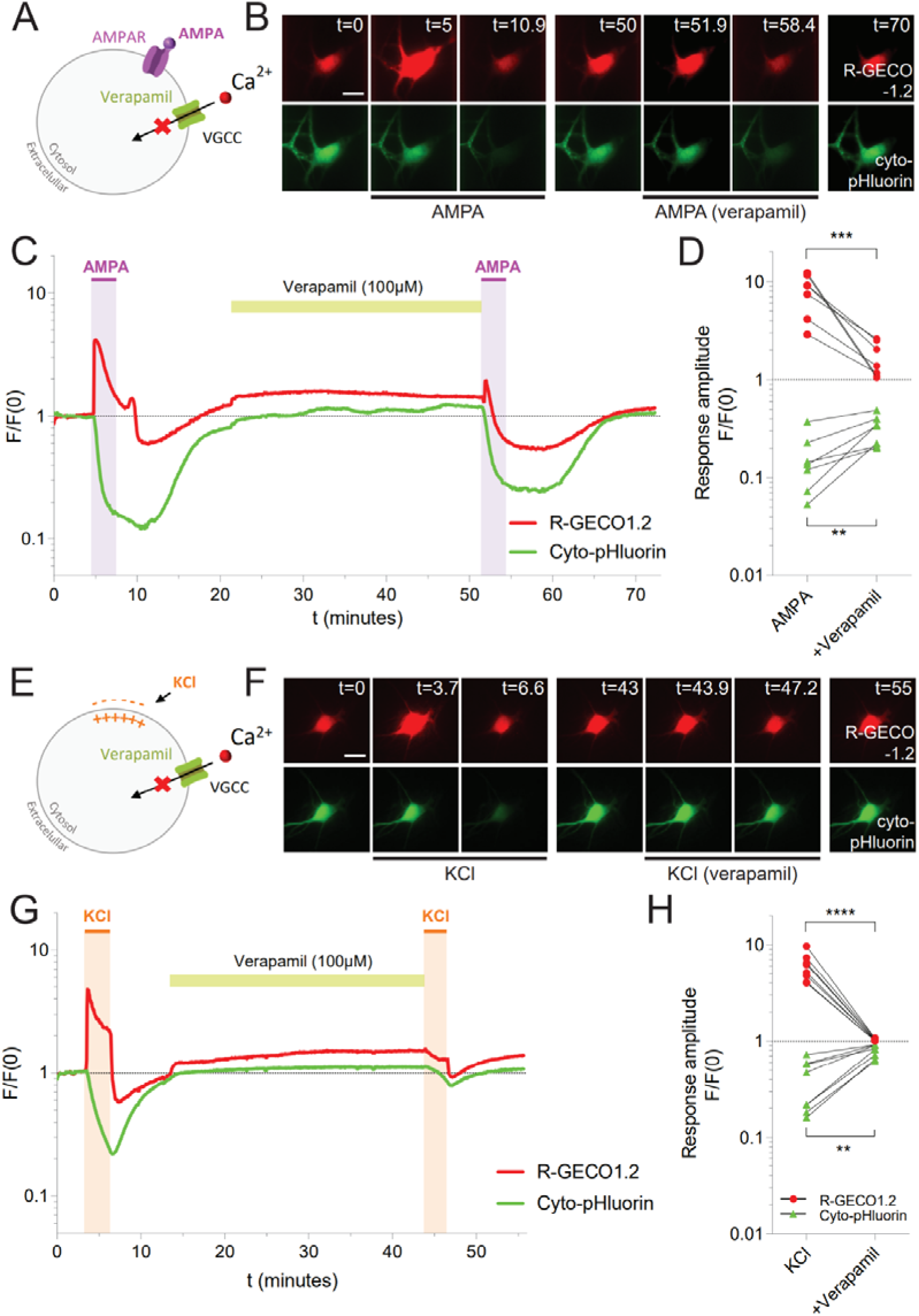
Activation of voltage-gated Ca^2+^-channels contribute to both AMPA- and KCl-induced cytosolic acidification. *(A,E)* Schematic representation of the expected mechanism of action of Verapamil in the two different stimulation paradigms (*B, F*) Representative image sequences of hippocampal neurons co-expressing R-GECO1.2 (red) and cyto-pHluorin (green). (Scale bar, 20 μm). Images are plotted for the indicated time points and show the change in fluorescence in response to AMPA or KCl before and after addition of the voltage-gated Ca^2+^ channel inhibitor verapamil. (C, G) Representative time courses of the normalized fluorescence (F/F0) for the neurons shown in (B, F), plotted on a log10-scale. AMPA (100 µM) or high KCl (60 mM) was applied at the indicated time points for 3 min before and after incubation with verapamil (100 µM) for 30 min. (*D, H*) Individual maximal responses induced by AMPA or KCl before and after verapamil incubation, shown on a log10-scale. Data are evaluated by ratio paired *t* tests, **P≤0.01, ***P≤0.001 ****P≤0.0001. AMPA (N = 7 cells), KCl (N = 8 cells). AMPA data include responses in presence of APV.

In KCl experiments, verapamil effectively prevented R-GECO1.2 fluorescence rise, indicating an effective inhibition of Ca^2+^ influx through voltage-gated Ca^2+^ channels (Fig.7E-H). Verapamil concomitantly reduced the maximal cyto-pHluorin responses, indicating reduced cytosolic acidification. This result is in line with a previous report, showing that depolarization-induced extracellular alkalization is reduced by nifedipine, a blocker of L-type Ca^2+^ channels (Makani and Chesler, 2010).

These results show that both AMPA- and KCl-induced cytosolic acidification are indeed dependent on voltage-gated Ca^2+^ channels to some degree. Verapamil left a significantly higher acidification intact following AMPA receptor stimulation than after KCl treatment (log(pHluorin_AMPA_) ± SEM = −0.524 ± 0.057 compared to log(pHluorin_KCl_) = −0.105 ± 0.024, Student t-test p = <0.0001). However, verapamil was also less efficient in reducing AMPA mediated Ca²⁺-influx (log(R-GECO1.2_AMPA_) ± SEM = 0.200 ± 0.065 compared to log(R-GECO1.2_KCl_) = 0.018 ± 0.003, Student t-test p = 0.0102), and consequently it is unclear how the residual acidification relates to the residual Ca²⁺-influx.

To further address a possible difference in Ca^2+^ dependence between AMPA- and KCl-induced acidification, we aimed to suppress cytosolic Ca^2+^ rise by preincubating hippocampal cultures with the membrane permeable Ca^2+^ chelator BAPTA-AM (Fig.8). Hippocampal neurons co-expressing cyto-pHluorin and R-GECO1.2 were pre-incubated with BAPTA-AM (20 µM) for 15-20 min prior to stimulation with AMPA or KCl (Fig.8A-F, G-L). BAPTA-AM treatment greatly reduced AMPA-induced maximal R-GECO1.2 responses compared to the previously obtained responses in the absence of BAPTA-AM (Fig.8D), indicating effective chelation of cytosolic Ca^2+^. In contrast, there was no significant difference in the AMPA-induced maximal cyto-pHluorin responses obtained in the presence and absence of BAPTA-AM, suggesting that the cytosolic acidification was not correspondingly reduced by the cytosolic Ca^2+^ chelation. Further analyses of the mean ratios between AMPA-induced cyto-pHluorin and R-GECO1.2 fluorescence responses did, however, not reveal any effect of BAPTA-AM (Fig.8E-F). In contrast, BAPTA-AM treatment completely abolished both R-GECO1.2 and cyto-pHluorin fluorescence responses induced by KCl (Fig.8J), and no difference could be detected in the response ratios (Fig.8K-L). Together, these results indicate that cytosolic Ca^2+^ chelation efficiently eliminates KCl-induced cytosolic acidification, suggesting complete Ca^2+^ dependence, whereas AMPA-induced cytosolic acidification appears to include a Ca²⁺-independent component.

**Figure 8.**
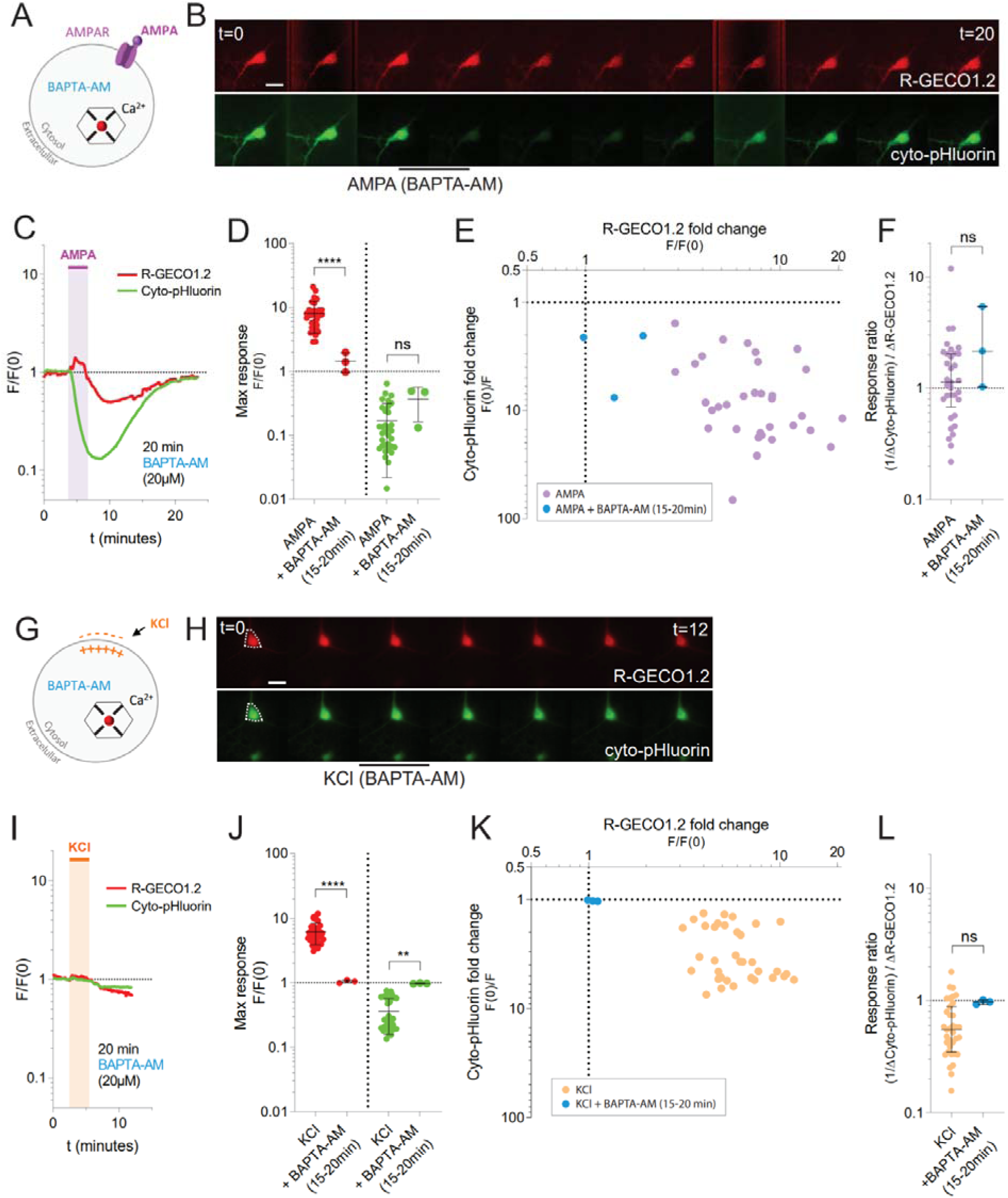
Chelation of intracellular Ca^2+^ prevents KCl-induced cytosolic acidification but does not significantly reduce acidification from AMPA. (*A,G*) Schematic representation of Ca^2+^ chelation by BAPTA-AM in the two different stimulation paradigms (*B, H*) Representative image sequences of a hippocampal neuron co-expressing R-GECO1.2 (red) and cyto-pHluorin (green). (Scale bar, 20 μm). Images are plotted for every two minutes of recording in the indicated time intervals and show the change in fluorescence in response to AMPA or KCl after pre-incubation with the Ca^2+^ chelator BAPTA-AM. (*C, I*) Representative time courses of the normalized fluorescence (F/F0) in the somatic region of the neurons shown in (A, F), plotted on log10-scales. AMPA (100 µM) or high KCl (60 mM) was applied at the indicated time points for 3 min, after pre-incubation with BAPTA-AM (20 µM) for 20 min. (*D, J*) Comparison of the maximal responses induced by AMPA or KCl in presence of BAPTA-AM (15-20 min pre-incubation) compared to the maximal responses induced by AMPA or KCl in absence of BAPTA-AM. The maximal response values reflect the maximal fold change in fluorescence for R-GECO1.2 and cyto-pHluorin in individual experiments. Data are shown on a log10-scale with mean ± SD. Log-transformed R-GECO1.2 or cyto-pHluorin responses are evaluated by unpaired t-tests, ***P≤0.001, ****P≤0.0001. (*E, K*) x-y plot showing the maximal cyto-pHluorin response as a function of the maximal R-GECO1.2 response for individual cells treated with AMPA or KCl in presence of BAPTA-AM (15-20 min pre-incubation) compared to the maximal responses induced by AMPA or KCl in absence of BAPTA-AM. The maximal response values reflect the maximal fold change in fluorescence for R-GECO1.2 and cyto-pHluorin in individual experiments, with cyto-pHluorin responses plotted as inversed values (F(0)/F). Data are shown on log10-scales. (*F, L*) Comparison of cyto-pHluorin to R-GECO1.2 maximal response ratios for individual experiments shown in (E, K). Ratios are calculated with inversed values of cyto-pHluorin responses (F(0)/F), and are shown on a log10-scale with medians and interquartile ranges. Data are evaluated by Mann-Whitney tests, ns: P>0.1. AMPA (N = 3 cells), KCl (N = 3 cells).

### Acidification induced by AMPA, but not simple depolarization, relies on a plasma membrane proton gradient

To investigate possible Ca²⁺-independent components of receptor and depolarization induced cytosolic acidification, we next assessed the requirement for an inward proton gradient across the plasma membrane for AMPA- and KCl-induced pH shifts, as a potential source of H^+^. It has previously been shown, using hippocampal slices, that neuronal surface pH remains alkaline at membrane depolarization to voltages positive to the equilibrium potential for H^+^ (Makani and Chesler, 2010). Based on this finding, depolarization-induced neuronal pH shifts are expected to depend on active transport and not a passive flux of H^+^ (or a base equivalent) through a voltage-dependent ion channel. Therefore, we wanted to assess whether protons could be conducted through the plasma membrane and if this differs between AMPA- and KCl-induced cytosolic acidification.

Initially, the extracellular pH level required to reach reversal potential for protons (V_rev,H+_) across the plasma membrane was determined in dissociated hippocampal neurons co-expressing cyto-pHluorin and R-GECO1.2 (Fig.9A-D). No change in baseline fluorescence was observed during a gradual decrease of extracellular pH from 9.0 to 7.4 (pH 9.0, 8.5, 8.0, 7.4) at resting conditions (Fig.9A-B). When a protonophore, CCCP (5 µM), was added during the gradual pH decrease the cyto-pHluorin fluorescence remained stable at extracellular pH 9.0, but decreased step-wise at the gradual decrease from pH 8.5 to 7.4 (Fig.9C-D). This indicated a neutral electrochemical H^+^ gradient across the plasma membrane at extracellular pH ~9.0, which is in agreement with theoretical calculations. We then adjusted the extracellular buffer pH to 9 and administered AMPA or KCl (3 min) to dissociated hippocampal neurons co-expressing cyto-pHluorin and R-GECO1.2 (Fig.9E-G,K-M). At extracellular pH 9, AMPA stimulations induced maximal R-GECO1.2 fluorescence responses that were comparable to the responses acquired at pH 7.4 (Fig.9H). In contrast, maximal cyto-pHluorin fluorescence responses were significantly lower compared to responses at pH 7.4. Moreover, a comparison of the fluorescence response ratios (Fig.9I-J) showed a significant decrease in the cyto-pHluorin to R-GECO1.2 ratio, indicating that Ca^2+^ rises were associated with lower cytosolic acidification at pH 9. Together, these data show that AMPA-induced cytosolic acidification is reduced at extracellular pH 9 regardless of any shift in intracellular Ca^2+^, indicating a H^+^-gradient dependent component of AMPA receptor-induced acidification.

**Figure 9.**
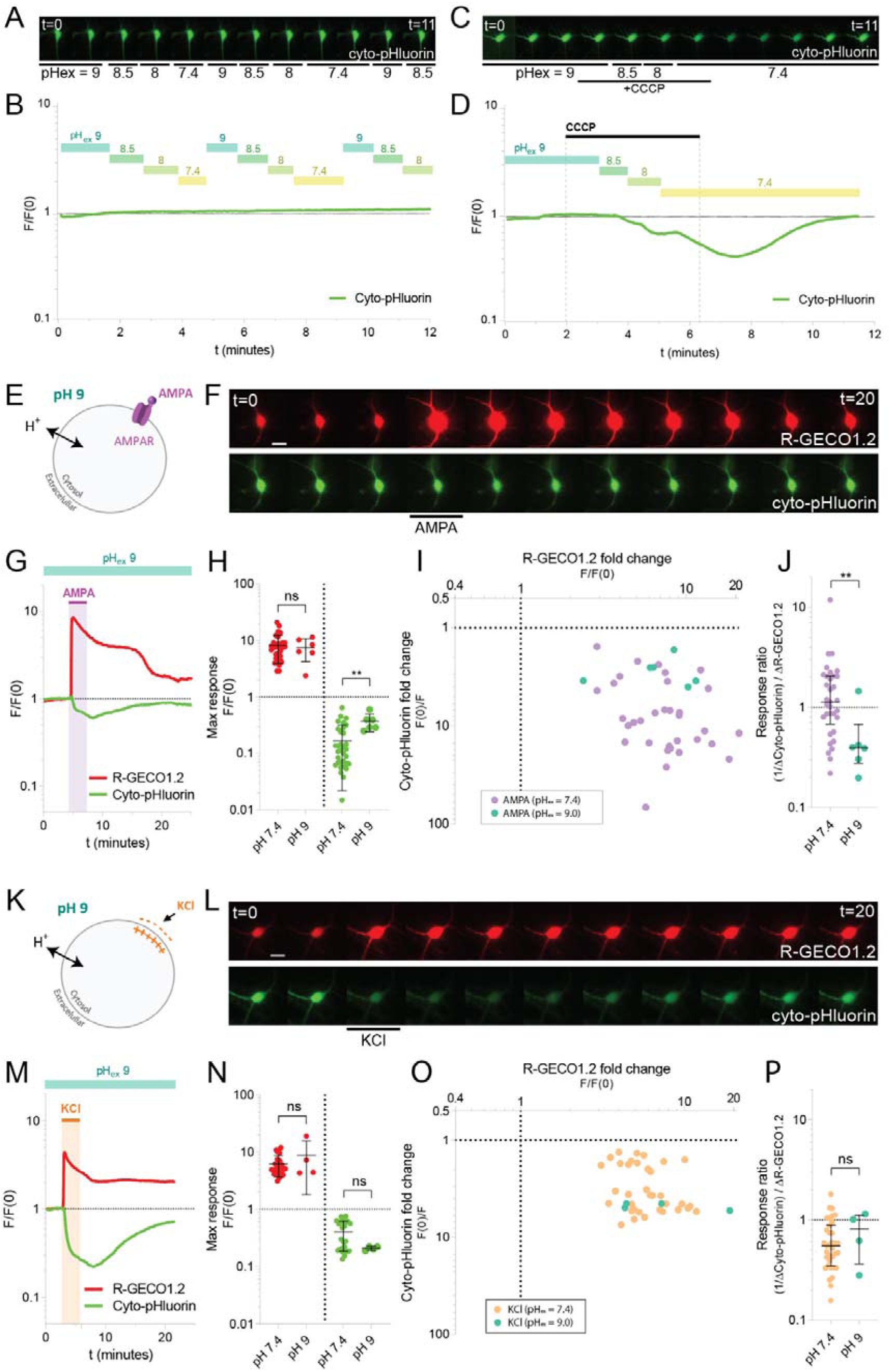
Elevation of extracellular pH to diminish the inward proton gradient reduces AMPA-induced acidification but does not significantly reduce acidification from KCl. (*A, B*) Representative image sequence and time course of the normalized fluorescence for a hippocampal neuron expressing cyto-pHluorin (green). Images are plotted for every minute of recording from 0-11 min and show the cell fluorescence during the gradual decrease of extracellular pH from 9.0 to 7.4 (pH 9.0, 8.5, 8.0, 7.4). The time course shows the normalized fluorescence (F/F0) in the somatic region of the cell, plotted on a log10-scale. The extracellular buffers were each applied for 1 min at the indicated time points. N = 1 cell. (*C, D*) Representative image sequence and time course of the normalized fluorescence for a hippocampal neuron expressing cyto-pHluorin (green). Images are plotted for every minute of recording from 0-11 min and show the cell fluorescence during the gradual decrease of extracellular pH from 9.0 to 7.4 (pH 9.0, 8.5, 8.0, 7.4) in presence of the protonophore CCCP (5 µM). The time course shows the normalized fluorescence (F/F0) in the somatic region of the cell, plotted on a log10-scale. The extracellular buffers were each applied for 1 min in presence of CCCP. N = 5 cells. (*E,K*) Schematic representation of the influence of extracellular pH 9 in the proton gradient in the two different stimulation paradigms (*F, L*) Representative image sequences of a hippocampal neuron co-expressing R-GECO1.2 (red) and cyto-pHluorin (green). (Scale bar, 20 μm). Images are plotted for every two minutes of recording from 0 to 20 min and show the change in fluorescence in response to AMPA or KCl at extracellular pH 9. (*G, M*) Representative time courses of the normalized fluorescence (F/F0) in the somatic region of the neurons shown in (F, L), plotted on log10-scales. AMPA (100 µM) or KCl (60 mM) was applied at the indicated time points for 3 min in extracellular buffers adjusted to pH 9. (*H, N*) Comparison of the maximal responses induced by AMPA or KCl at extracellular pH 9 compared to the maximal responses induced by AMPA or KCl at extracellular pH 7.4. The maximal response values reflect the maximal fold change in fluorescence for R-GECO1.2 and cyto-pHluorin in individual experiments. Data are shown on a log10-scale with mean ± SD. Log-transformed R-GECO1.2 or cyto-pHluorin responses are evaluated by unpaired t-test, **P≤0.01. (*I, O*) x-y plot showing the maximal cyto-pHluorin response as a function of the maximal R-GECO1.2 response for individual cells treated with AMPA or KCl in at extracellular pH 9 compared to the maximal responses induced by AMPA or KCl at extracellular pH 7.4. The maximal response values reflect the maximal fold change in fluorescence for R-GECO1.2 and cyto-pHluorin in individual experiments, with cyto-pHluorin responses plotted as inversed values (F(0)/F). Data are shown on log10-scales. (*J, P*) Comparison of cyto-pHluorin to R-GECO1.2 maximal response ratios for individual experiments shown in (H, M). Ratios are calculated with inversed values of cyto-pHluorin responses (F(0)/F), and are shown on a log10-scale with medians and interquartile ranges. Data are evaluated by Mann-Whitney tests, **P≤0.01. AMPA/pH9 (N = 6 cells), KCl/pH9 (N = 4 cells).

Conversely, KCl-induced cyto-pHluorin and R-GECO1.2 fluorescence responses were not significantly different at extracellular pH 9 compared to responses obtained at pH 7.4 (Fig.9N-P). This indicates that depolarization-induced cytosolic acidification is not dependent on an electrochemical driving force for H^+^ across the plasma membrane, which is in agreement with the previous report on depolarization-dependent extracellular alkalization from Makani and Chesler (2010).

## Discussion

Neuronal Ca^2+^ imaging is a well-established technique for monitoring neuronal activity, linking Ca^2+^ dynamics to synaptic plasticity and excitotoxicity. However, the relationship between neuronal activity and cytosolic pH shifts remains underexplored, partly due to technical limitations, such as a lack of specific inhibitors. While various neuronal types have been reported to exhibit activity-dependent intracellular pH shifts, most studies have focused on conditions of excitotoxicity, seizure-like activity, or the bath application of agonists. This has left the physiological intracellular pH dynamics associated with normal neuronal activity largely understudied. Furthermore, activity-dependent pH fluctuations have been linked to Ca^2+^, however, the complex interplay between Ca^2+^ and pH has been challenging to investigate comprehensively, as studies often measure these systems separately, which can lead to misinterpretations. Here, using a dual biosensor approach, we simultaneously measured cytosolic Ca^2+^ and pH in hippocampal neurons under LTP and pharmacological stimulations to address 1) physiological intracellular pH dynamics and 2) its relationship to Ca^2+^ during neuronal activity.

Prior studies have focused on interstitial pH changes in the CA1 region following Schaffer collateral stimulation, where alkaline transients have been reported (Krishtal et al., 1987; Chen and Chesler, 1992) that enhance NMDA receptor responses (Makani and Chesler, 2007). However, our dual-sensor approach provides the first evidence of single-cell intracellular pH shifts recorded simultaneously with Ca^2+^ responses in CA1 neurons during LTP-inducing TBS. This finding, along with our measurements of spontaneous activity, suggests that cytosolic pH shifts are a characteristic of physiological neurotransmission, not just during seizure-like conditions as previously reported. The potential for cytosolic H^+^ ions to act as general modulators of physiological neurotransmission has been suggested (Sinning and Hübner, 2013) however, the theory is still not well-established. Studies do indicate that general intracellular acidification, or inhibited acid extrusion, can decrease synaptic activity in hippocampal slices (Lee et al., 1996; Bonnet et al., 2000; Xiong et al., 2000) and increase the seizure threshold in *in vivo* models of epilepsy (Sinning et al., 2011). Cytosolic acidification has also been shown to ameliorate Parkinson’s disease (Komilova, 2021) and obsessive-compulsive disorder by decreasing firing activity (Hatamaka, 2024), highlighting the neuroprotective effects of activity-induced acid shifts. Furthermore, activity-induced cytosolic acidification has been linked to other synaptic processes; it induces spontaneous, synchronous zinc spikes in hippocampal neurons, which are essential for synapse maturation and neuronal differentiation (Chen Zhang, 2021). It also increases RNA editing by adenosine deaminase acting on RNA (ADAR) proteins, with overexpression or knockdown of these proteins shown to decrease or increase neuronal excitability, respectively (Malik, 2021). Therefore, given the implications of intracellular pH in neuronal signaling, activity-induced cytosolic acid transients may play a more active role in physiological neurotransmission than previously predicted, acting as a secondary messenger.

One of the factors proposed to contribute to neuronal acidification upon activity is the generation of glycolytic byproducts. It has been reported that the replacement of glucose with pyruvate, to bypass glycolysis, caused a great reduction in NMDA-induced neuronal acidification in hippocampal slice cultures, supposedly due to decelerated glycolysis (Zhan et al., 1997, 1998). However, here we found that pyruvate substitution completely prevented both Ca^2+^ and pH responses when specifically induced by NMDA. Our results in *Xenopus* oocytes further showed that this is due to a direct inhibitory effect of pyruvate on the NMDA receptors. Pyruvate has been reported to provide neuroprotection against excitotoxicity *in vitro* and *in vivo* through various mechanisms (Izumi and Olney, 1995; Wang et al., 2009; Izumi and Zorumski, 2010; Ryou et al., 2012; Tian et al., 2014). Thus, this direct inhibition poses a novel mechanism of pyruvate-mediated neuroprotection.

Our data from hippocampal cultures revealed significant cell-to-cell variability in biosensor response sizes under different pharmacological stimulations. By using a dual sensor approach, we assessed Ca²⁺ and pH responses in individual cells. Although we found no direct correlation between their maximal responses for any of the stimulation types, a higher Ca²⁺ influx in a given neuron elicited a stronger pH response with a constant maximal response ratio (H⁺/Ca²⁺). In contrast, the median maximal response ratios did vary significantly between NMDA, AMPA, and KCl stimuli, indicating distinct stimulus-specific response patterns and potential differential mechanisms. Previous studies have reported a clear Ca²⁺ dependency of the cytosolic acidification induced by both membrane depolarization and pharmacological glutamate receptor stimulation. Consistent with these findings, our data show that depolarization-induced cytosolic acidification strongly depends on voltage-dependent Ca²⁺ channels and a rise in cytosolic Ca²⁺, supporting the hypothesis that such pH shifts involve active proton import (Makani and Chesler, 2010; Cheng et al., 2008). In two previous studies, KCl-mediated depolarization of hippocampal neurons also induced cytosolic alkalization (linked to extracellular Ca^2+^) that limited acid transients (Cheng et al., 2008; Svichar et al., 2011). The fluorescence intensity of cyto-pHluorin is less pH sensitive above pH 7.4, and consequently, we would not expect to detect these alkaline transients in our setup. Remarkably, we uncovered a Ca²⁺-independent acid component during glutamate receptor stimulation that has not been previously described; acidification following AMPA receptor stimulation does not exclusively require cytosolic Ca²⁺ but also relies on the plasma membrane proton gradient, suggesting that AMPA receptors may facilitate passive proton influx that amplifies the acid response. Since NMDA treatments were also associated with large acid shifts, a similar mechanism might underlie NMDA receptor-mediated pH changes. However, this is more experimentally challenging to assess due to the pH sensitivity of the NMDA receptor (Ladislav et al., 1990). Further investigation is required to determine if ionotropic glutamate receptors have inherent proton permeability, and if so, whether this proton flux is functionally relevant.

In summary, these data provide the first evidence of simultaneous cytosolic Ca^2+^ and pH shifts during neurotransmission. They additionally reveal the heterogeneity of activity-induced acidification transients, both in intensity and mechanism, depending on the type of stimulation. Notably, we identify a new Ca²⁺-independent component of the acidification and reevaluate glycolysis as a contributing factor. Based on our results, we propose that pH dynamics could serve as a proxy for neuronal activity, and that cytosolic acid transients induced by activity may act as secondary messengers with potentially distinct signaling roles depending on the neurotransmission type. The future development of techniques to specifically assess and modulate pH dynamics in hippocampal neuron microdomains—such as presynaptic terminals, dendritic spines, and postsynaptic densities—will be crucial for understanding endogenous pH fluctuations and their potential implications for synaptic plasticity.

## Acknowledgements

We thank Nabeela Khadim, Lone Rosenquist, and Donna G. Czerny for their excellent technical assistance. We also thank Prof. Hans Bräuner-Osborne (University of Copenhagen, Denmark) and Ass. Prof. Kasper B. Hansen (University of Montana, Missoula, USA) for insightful comments on glutamate receptor pharmacology and the expression of recombinant receptors in *Xenopus* oocytes. A special thanks to M.Sc. Søren Emil Nørr (University of Copenhagen, Denmark) for his assistance in producing the cyto-pHluorin AAVs.

The work was supported by the Lundbeck Foundation (R191-2015-1385) (SEP, KLM) and the Danish Council for Independent Research - Medical Sciences (4004-00555B) (KLM) and (DFF–1333-00142) (TK).

## Footnotes

1 It should be noted that different effective concentrations of glutamate receptor agonists were used (EC_20_ NMDA vs. saturating AMPA concentration), and smaller NMDA responses likely reflect submaximal receptor activation. The NMDA stimulations were based on the conventional chemical LTD protocol (Lee et al., 1998).

## Notes

### Competing Interest Statement

The authors have declared no competing interest.

