## supplementary figures for "Glutamate receptor-dependent cytosolic acidification in hippocampal neurons involves passive flux of protons from the extracellular space"

Running title: Activity-induced acidification in hippocampal neurons

Sofie E. Pedersen^1*^, Ainoa Konomi-Pilkati^1*^, Nikolaj W. Hansen^2^, Trine Kvist^3^, Leonie P. Posselt^1^, Thorvald F. Andreassen^1^, Andreas T. Sørensen^1^, Jean François Perrier^2^, Kenneth L. Madsen^1†^

Supplementary figures

**S1.**
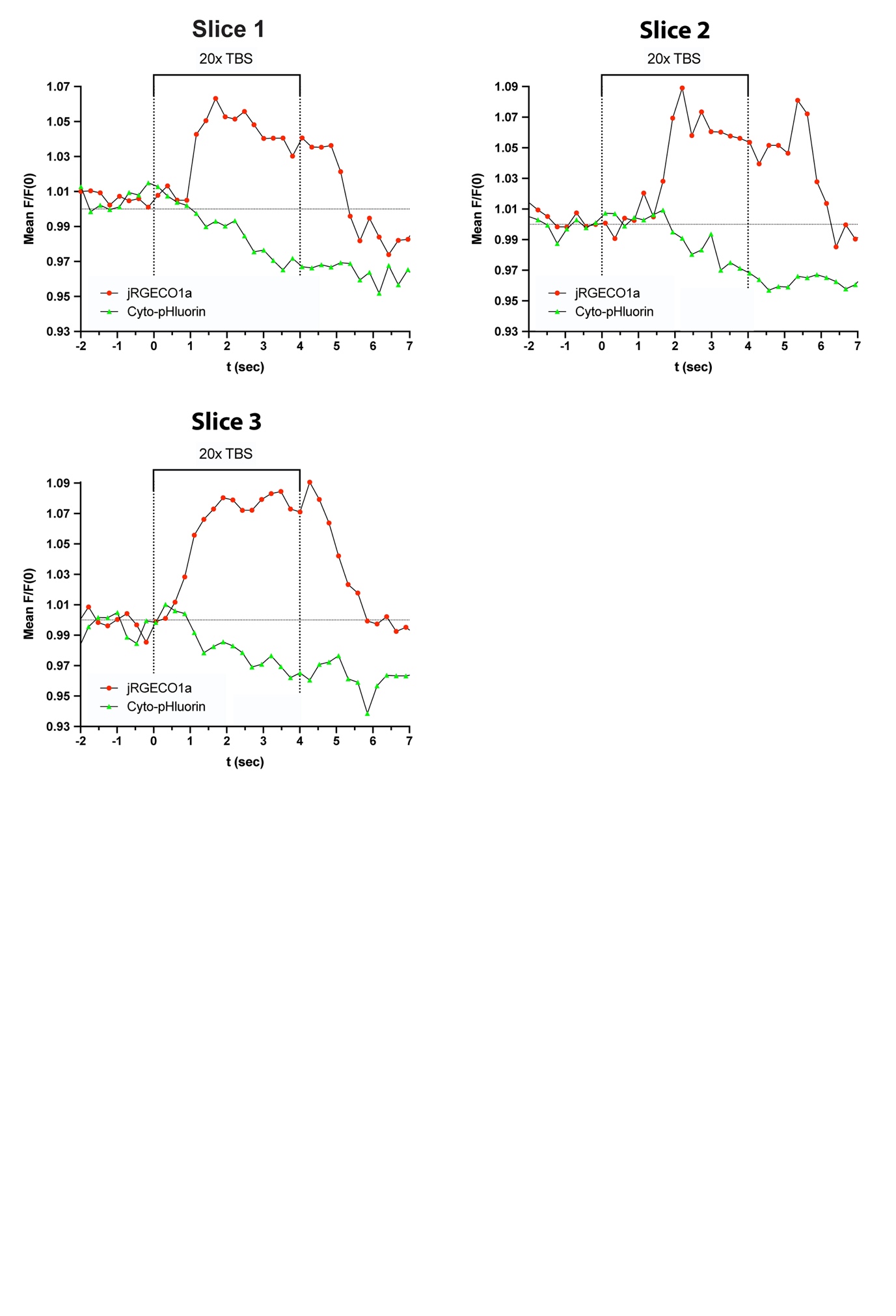

**Supplementary figure 1** *Response initiation time for cyto-pHluorin and jRGECO1a in hippocampal slices.*

The graphs show the normalized mean fluorescence (mean F/F0) for jRGECO1a (red) and cyto-pHluorin (green) recorded from the CA1 somas during the first seconds of LTP-inducing stimulation (20x TBS) at the Schaffer collaterals. The data were obtained in three independent slice experiments. Slice 1 represents the experiment shown in figure 1 C-E.

**S2.**

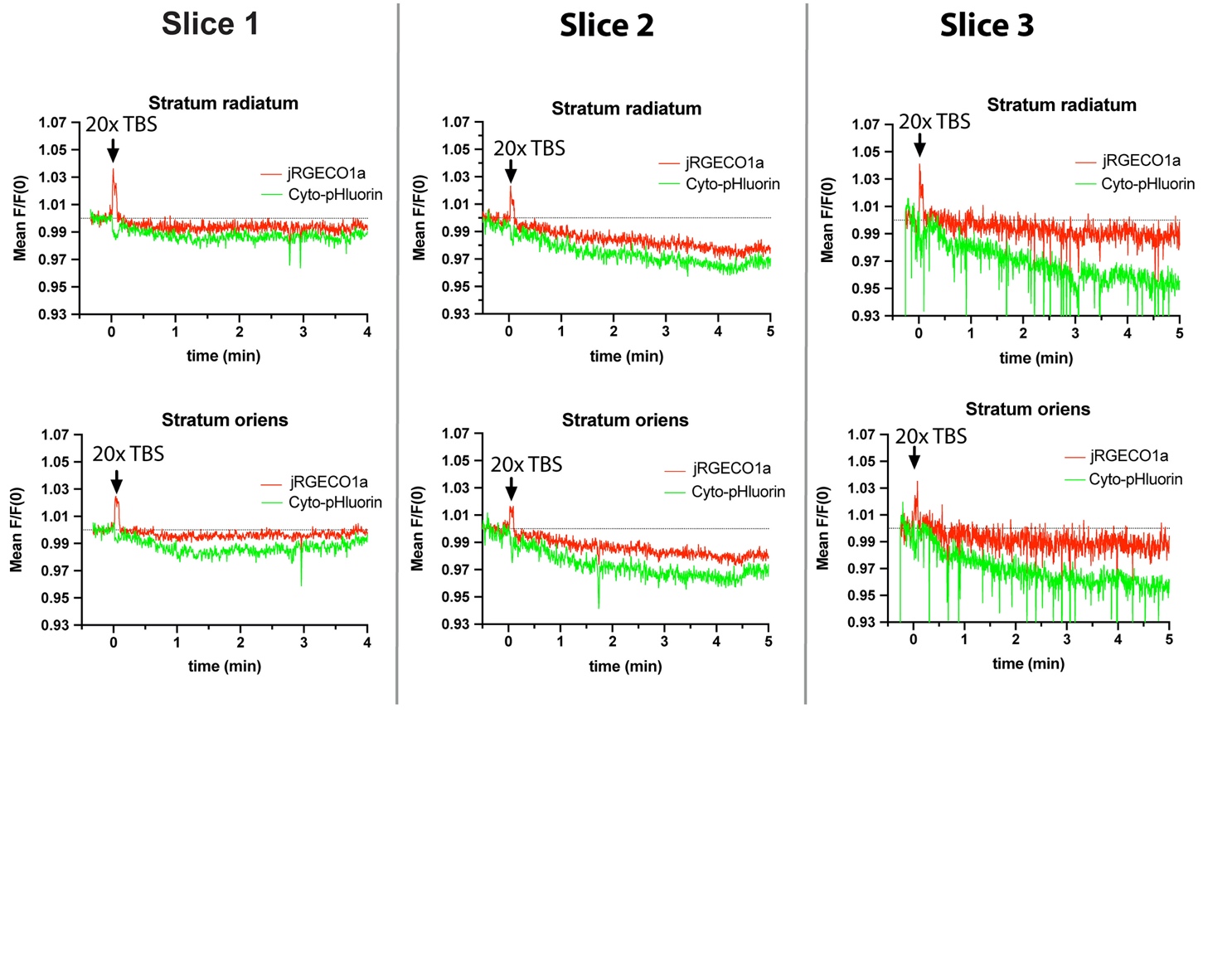

**Supplementary figure 2** *Fluorescence responses recorded from the CA1 stratum radiatum and oriens*.

The time courses of the mean normalized fluorescence (mean F/F0) for jRGECO1a (red) and cyto-pHluorin (green) in CA1 stratum radiatum and stratum oriens before, during and after LTP-inducing stimulation (20x TBS) at the Schaffer collaterals. The data were obtained in three independent slice experiments. Slice 1 represents the experiment shown in figure 1 C-E.

**S3.**

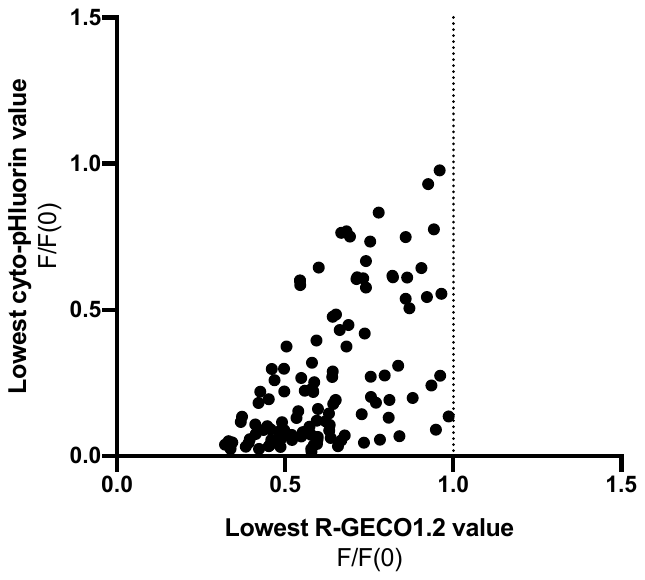

**S4.**

**Supplementary figure 3** *The extent of R-GECO1.2 downward shifts after treatments correlates with the extent of cytosolic acidification.*

The extent of the downward R-GECO1.2 fluorescence shift, represented by the lowest R-GECO1.2 value achieved after a stimulation, plotted as a function of the maximal cyto-pHluorin response. Data for all treatments (glutamate, NMDA, AMPA and KCl) eliciting a R-GECO1.2 fluorescence response with a downward shift to below baseline level are included. Pearson correlation coefficient, ****P <0.0001.

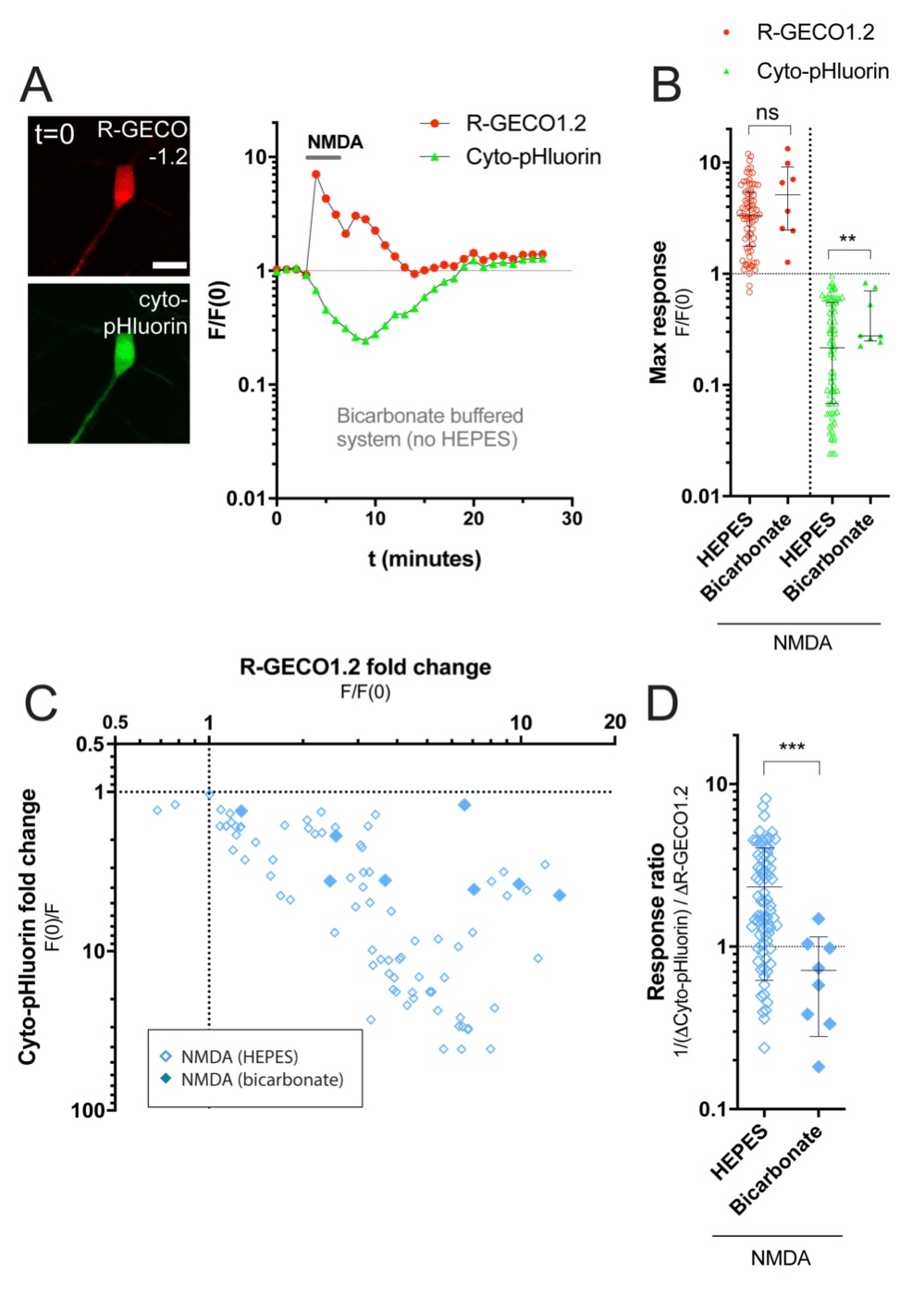

**Supplementary figure 4** *A bicarbonate-based buffer system reduces, but does not prevent, NMDA-induced acidification.*

*(A*) Representative images and representative time course of the normalized fluorescence (F/F_0_) for a hippocampal neuron co-expressing R-GECO1.2 (red) and cyto-pHluorin (green). (Scale bar, 20 μm). NMDA (20 µM NMDA, 10 µM glycine) was applied for 3 min in a HEPES free bicarbonate buffered system. Data are shown on a log10-scale. (*B*) Comparison of the maximal NMDA responses induced in bicarbonate and HEPES buffered medium. The maximal response values reflect the maximal fold change in fluorescence for R-GECO1.2 and cyto-pHluorin in individual experiments. Data are shown on a log10-scale with mean ± SD. Log-transformed R-GECO1.2 or cyto-pHluorin responses are evaluated by unpaired t-tests, **P≤0.01. (*C*) x-y plot showing the maximal cyto-pHluorin response as a function of the maximal R-GECO1.2 response for individual cells in response to NMDA in bicarbonate or HEPES buffered medium. The maximal response values reflect the maximal fold change in fluorescence for R-GECO1.2 and cyto-pHluorin in individual experiments, with cyto-pHluorin responses plotted as inversed values (F(0)/F). Data are shown on log10-scales. (*D*) Comparison of cyto-pHluorin to R-GECO1.2 maximal response ratios for individual experiments shown in (C). Ratios are calculated with inversed values of cyto-pHluorin responses (F(0)/F), and are shown on a log10-scale with medians and interquartile ranges. Data are evaluated by a Mann-Whitney test, ***P≤0.001. Bicarbonate data (N = 8 cells).

**S5.**

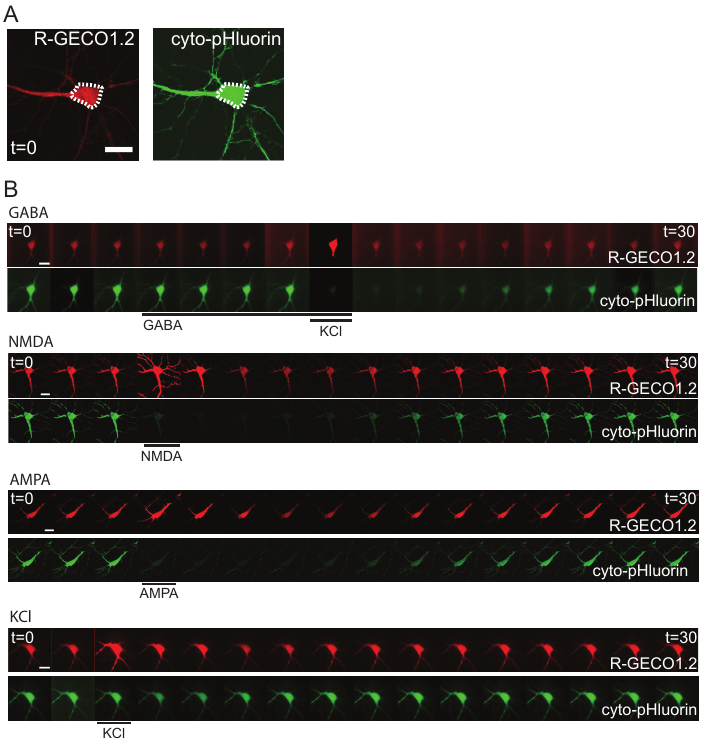

**Supplementary figure 5** *Image sequences used for fluorescence analyses in figure 2D-G.*

Representative image sequences used for analyses of fluorescent time courses in figure 2D-G. Images are plotted for every two minutes of recording from 0 to 30 min. (Scale bar, 20 µm). NB: images collected from GABA experiments are shown with different gains, since the illumination varied during recordings. GABA response quantifications are based on the original data.

**S6.**

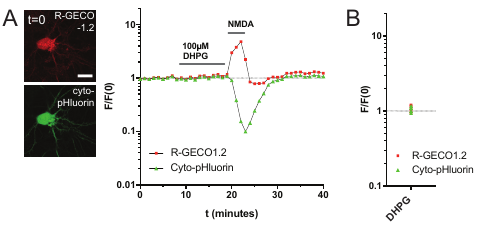

**Supplementary figure 6** *The group I metabotropic glutamate receptor (mGluR) agonist DHPG has no effect on R-GECO1.2 and cyto-pHluorin fluorescence in dissociated hippocampal neurons*.

(*A*) Representative image and time course of the normalized fluorescence (F/F0) for a hippocampal neuron, co-expressing R-GECO1.2 (red) and cyto-pHluorin (green) during DHPG stimulation. DHPG (100 µM) was applied at the indicated time point for 10 minutes, before control stimulating with NMDA (20 µM NMDA, 10 µM glycine) for 3 minutes. Both agonists were applied in buffer containing 2 mM MgCl. Data are plotted on a log10-scale. (*B*) Individual maximal responses induced by DHPG, shown on a log10-scale. N = 4 cells.

**S7.**

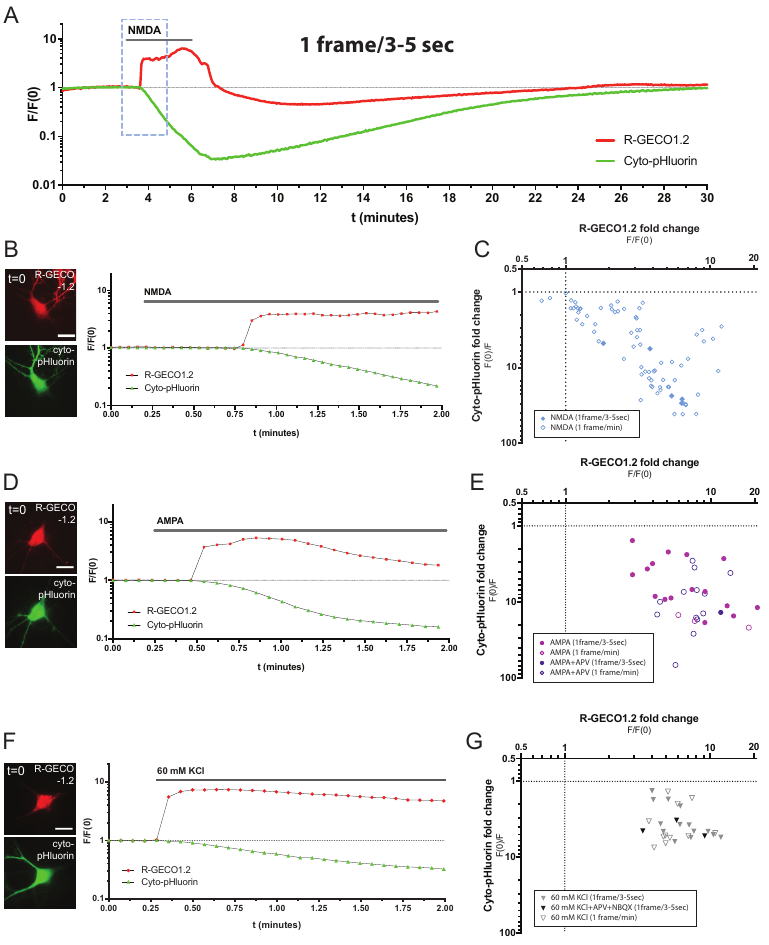

**Supplementary figure 7** *Response initiation interval assessed by fluorescent imaging with higher temporal resolution (image acquisition interval: 3-5 s).*

*(A*) Representative time course of the normalized fluorescence (F/F0) from R-GECO1.2 (red) and cyto-pHluorin (green), recorded from the somatic region of a hippocampal neuron with an acquisition rate of 1 image/3-5 s. NMDA (20 μM NMDA, 10 μM glycine) was applied at the indicated time point for 3 min. The broken line indicates the response initiation interval, which is shown in isolation in (B). (*B, D, F*) The initiation intervals of the fluorescence responses recorded from hippocampal neurons co-expressing R-GECO1.2 (red) and cyto-pHluorin (green) with an acquisition rate of 1 image/3-5 s. (Scale bar, 20 µm). NMDA (20 µM NMDA, 10 µM glycine), AMPA (100 µM) or KCl (60 mM) was applied at the indicated time point. (*C*, *E*) x-y plot for NMDA and AMPA data displayed in figure 2 I, showing the maximal fluorescence responses with specified image acquisition rates. Closed symbols: image acquisition interval of 3-5 s, open symbols: image acquisition interval of 1 min. Data are shown on log10-scales, with cyto-pHluorin responses plotted as inversed values (F(0)/F). (*G*) x-y plot for KCl data displayed in figure 2 I, showing the maximal fluorescence responses with specified image acquisition rates and inhibitor administration. Closed grey triangles: no inhibitor administration, image acquisition interval of 3-5 s. Closed black triangles: co-application of glutamate receptor inhibitors (100 µM APV, 50 µM NBQX), image acquisition interval of 3-5 s. Open triangles: image acquisition interval of 1 min. Data are shown on log10-scales, with cyto-pHluorin responses plotted as inversed values (F(0)/F).

**S8.**

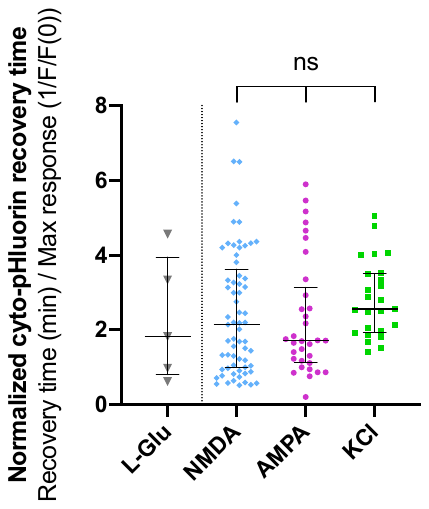

**Supplementary figure 8** *Cyto-pHluorin fluorescence recovery times do not differ significantly between treatments.*

For all treatments eliciting a cyto-pHluorin response (L-Glu, NMDA, AMPA, KCl), the cytosolic pH recovery time was estimated by calculating the interval between the time point of maximal cyto-pHluorin response until the time point of fluorescence recovery to baseline. To correct for differences in response size, the recovery time was subsequently normalized to the maximal cyto-pHluorin response value (1/(F/F(0)) in each experiment. Data from NMDA, AMPA and KCl experiments were statistically compared using Kruskal-Wallis and Dunn's multiple comparisons test, ns: P > 0.05.

**S9.**

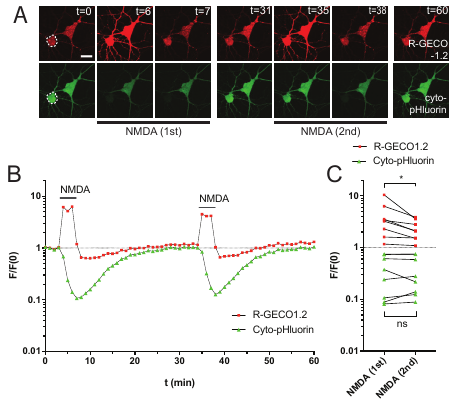

**Supplementary figure 9** *Repeated NMDA stimulations reduce the maximal response from R-GECO1.2 but has no significant effect on cyto-pHluorin responses*.

(*A*) Representative image sequence of hippocampal neuron co-expressing R-GECO1.2 (red) and cyto-pHluorin (green). (Scale bar, 20 µm). Images show the cell fluorescence at the indicated time points. The dotted line indicate the neuronal soma used in the analysis in (B). (*B*) Representative time course of the normalized fluorescence (F/F_0_) plotted on a log10-scale. NMDA (20 µM NMDA, 10 µM glycine) was applied for 3 min two consecutive times, allowing for recovery of baseline fluorescence between stimuli. (*C*) Individual maximal responses induced by the first NMDA stimulus (1^st^) and by the second NMDA stimulus (2^nd^), shown on a log10-scale. Data are evaluated by ratio paired *t* test, *P≤0.05. N = 8 cells.

**S10.**

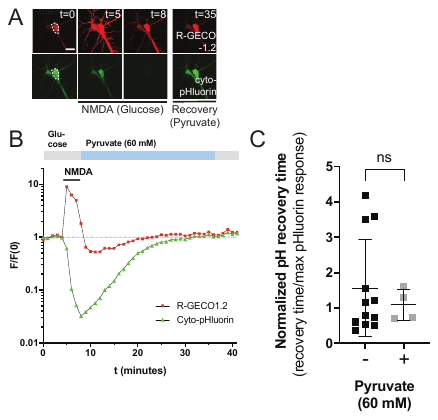

**Supplementary figure 10** *Post-stimulatory pyruvate substitution has no apparent effect on the recovery time of NMDA-induced acidification.*

(*A*) Representative image sequence of hippocampal neuron co-expressing R-GECO1.2 (red) and cyto-pHluorin (green). (Scale bar, 20 µm). Images show the cell fluorescence at the indicated time points. The dotted line indicates the neuronal soma used for the analyses in (B). *(B)* Representative time course of the normalized fluorescence (F/F_0_), plotted on a log10-scale. NMDA (20 µM NMDA, 10 µM glycine) was applied in conventional aCSF buffer at the indicated time point (3 min), before wash-out and fluorescence recovery in pyruvate-containing buffer. (*C*) Comparison of the cyto-pHluorin fluorescence recovery times displayed by neurons placed in glucose- or pyruvate-containing buffers immediately after NMDA stimulation. Only NMDA data acquired in the pyruvate experimental series (Fig. 2, S8, S9) were included in the analysis. The recovery time was defined as the interval from the maximal cyto-pHluorin response to the point of re-established baseline fluorescence. For each experiment, the recovery time was normalized to the inversed maximal fold change in cyto-pHluorin fluorescence (F(0)/F). Data are shown with mean ± SD and are evaluated by an unpaired *t*-test, ns: P>0.5. (–) pyruvate (N = 12 cells), (+) pyruvate (N = 4 cells.).

**S11.**

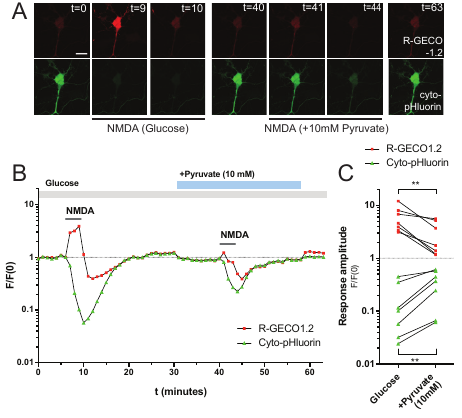

**Supplementary figure 11** *Addition of* *pyruvate (10 mM) to conventional aCSF buffer reduces NMDA-induced pH responses.*

*(A)* Representative image sequence of hippocampal neuron co-expressing R-GECO1.2 (red) and cyto-pHluorin (green). (Scale bar, 20 µm). Images show the cell fluorescence at the indicated time points. *(B)* Representative time course of the normalized fluorescence (F/F_0_) for the neuron in (A), plotted on a log10-scale. NMDA (20 µM NMDA, 10 µM glycine) was applied for 3 min before and after addition of 10 mM pyruvate to the extracellular medium. *(C)* Individual maximal fluorescence responses induced by NMDA in conventional aCSF before and after addition of pyruvate, shown on a log10-scale. Data are evaluated with ratio paired *t* test, **P≤0.01. N = 7 cells.

**S12.**

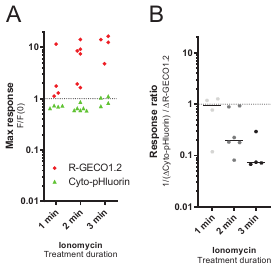

**Supplementary figure 12** *Maximal responses and response ratios for all ionomycin treatments shown in figure 5 E.*

(*A*) Individual maximal R-GECO1.2 (red) and cyto-pHluorin (green) fluorescence responses induced by ionomycin treatment for 1, 2 or 3 min, shown on a log10-scale. (*B*) Ratios of cyto-pHluorin to R-GECO1.2 maximal responses for individual experiments. Ratios are calculated with inversed values of cyto-pHluorin responses (F(0)/F), and are shown on a log10-scale. Ionomycin data are from 10 cells (N = 4-6 experiments for each treatment duration).

**S13.**

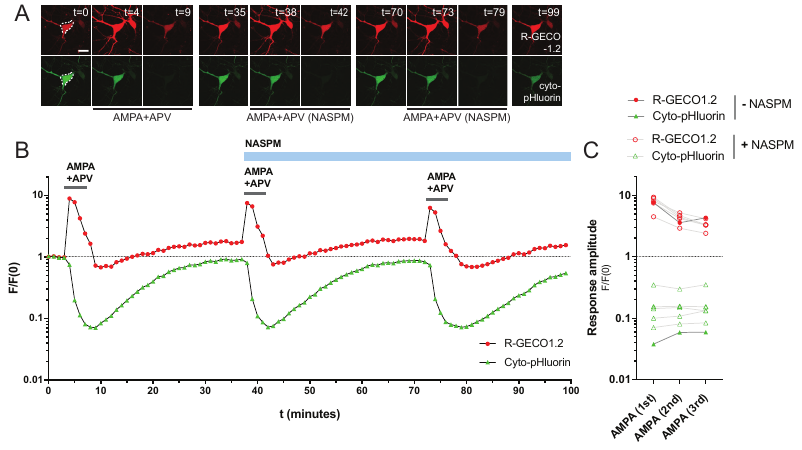

**Supplementary figure 13** *Ca^2+^-permeable- (CP-) AMPA receptors have no apparent effect on AMPA-induced cytosolic acidification.*

(*A*) Representative image sequence of a hippocampal neuron co-expressing R-GECO1.2 (red) and cyto-pHluorin (green). (Scale bar, 20 μm). Images are plotted for the indicated time points. The dotted line indicates the neuronal soma used in the analyses shown in (B). (*B*) Representative time course of the normalized fluorescence (F/F_0_), plotted on a log10-scale. AMPA (100 µM AMPA, 100 µM APV) was applied before and after addition of the specific CP-AMPA receptor inhibitor, NASPM (30 µM). (*C*) Individual maximal responses induced by consecutive AMPA stimulations in absence (closed symbols) or presence (open symbols) of NASPM during 2^nd^ and 3^rd^ stimulus. Data are shown on a log10-scale. + NASPM (N = 5 neurons), -NASPM (N = 1 neuron).

**Appendix 1:** *Fluorescence responses to TBS recorded in CA1 somas in three independent slice experiments (denoted slice 1-3)*.

A) Trace of the normalized field potential recordings before and after LTP induction through TBS stimulation. The inset displays representative fEPSP recordings before (black) and after (grey) LTP induction. B) The time course of the mean normalized fluorescence (mean F/F0) for jRGECO1a (red) and cyto-pHluorin (green) in CA1 somas measured from the hippocampal slices before, during and after LTP-inducing TBS at the Schaffer collaterals. C) The normalized fluorescence for the individual CA1 somas.

**Slice 1 (displayed in main figure 1 C-E)**

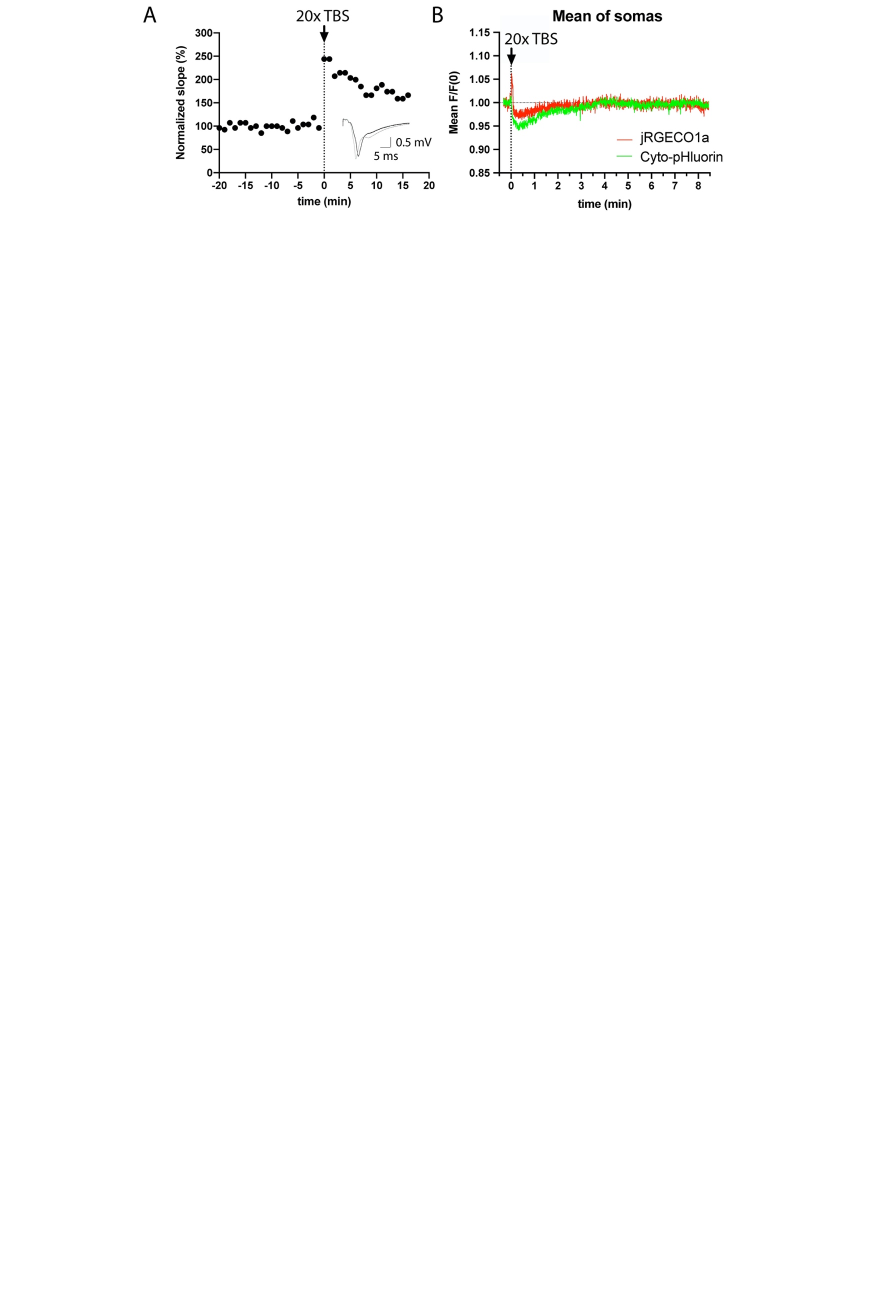

*(“Slice 1” continued on next page)*

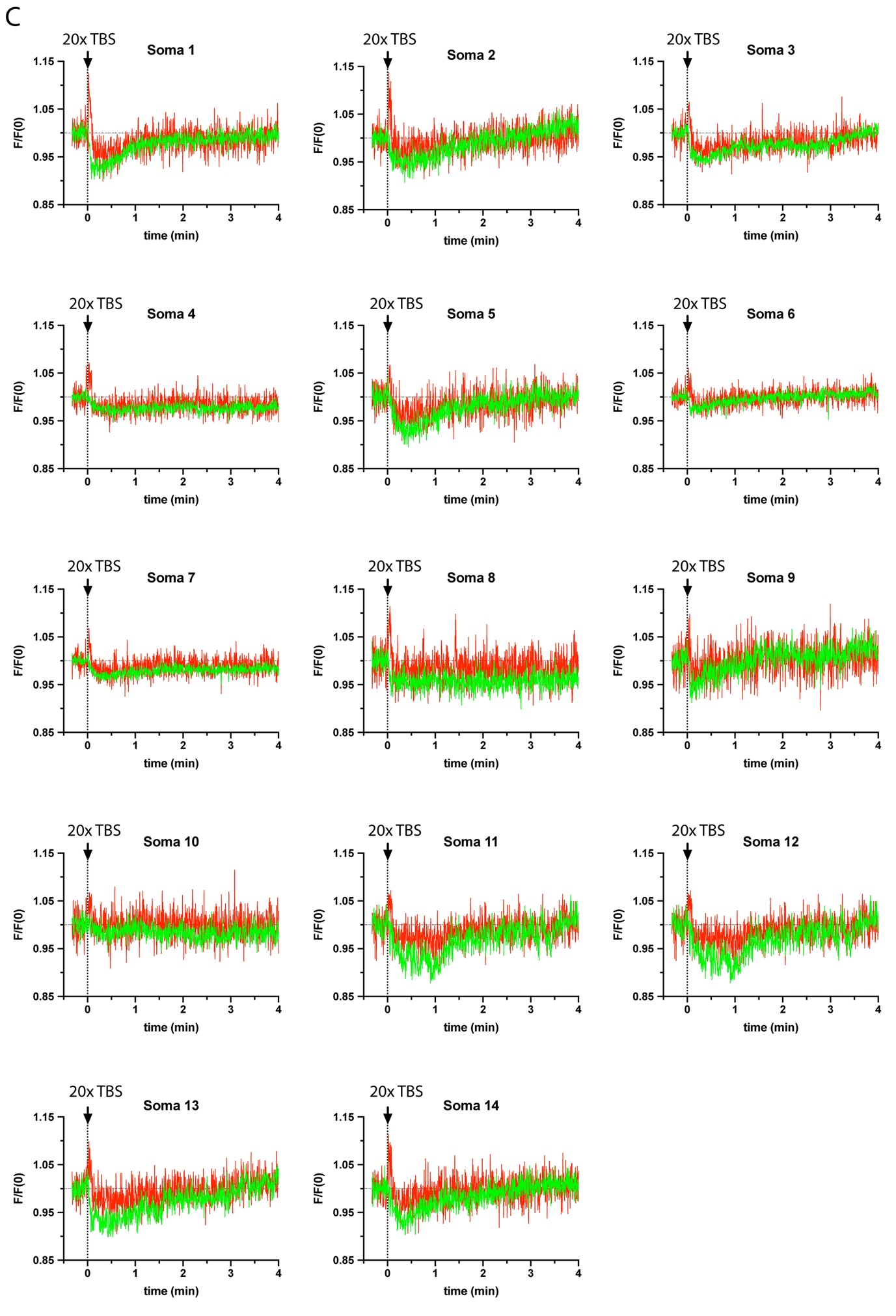

**Slice 2:**

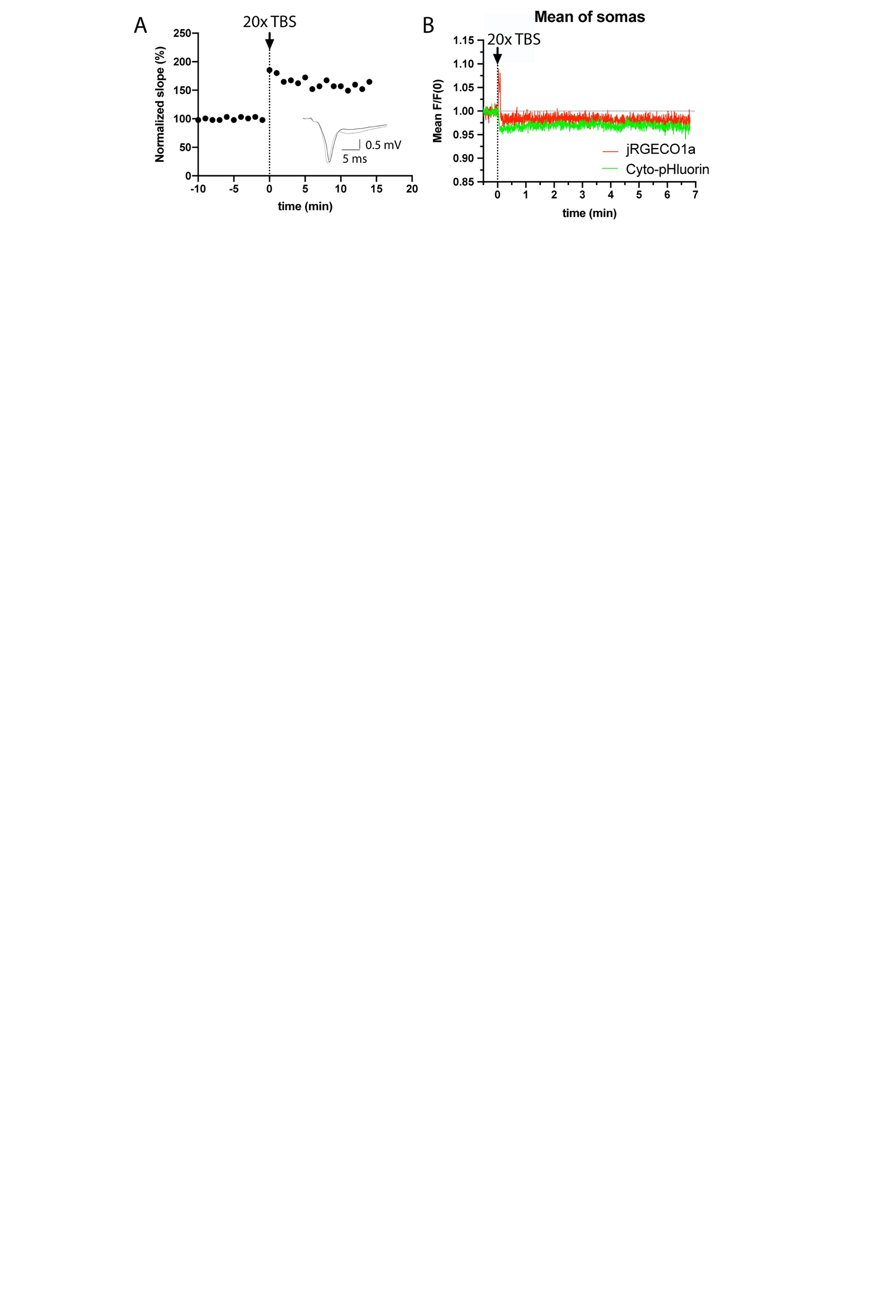

*(“Slice 2” continued on next page)*

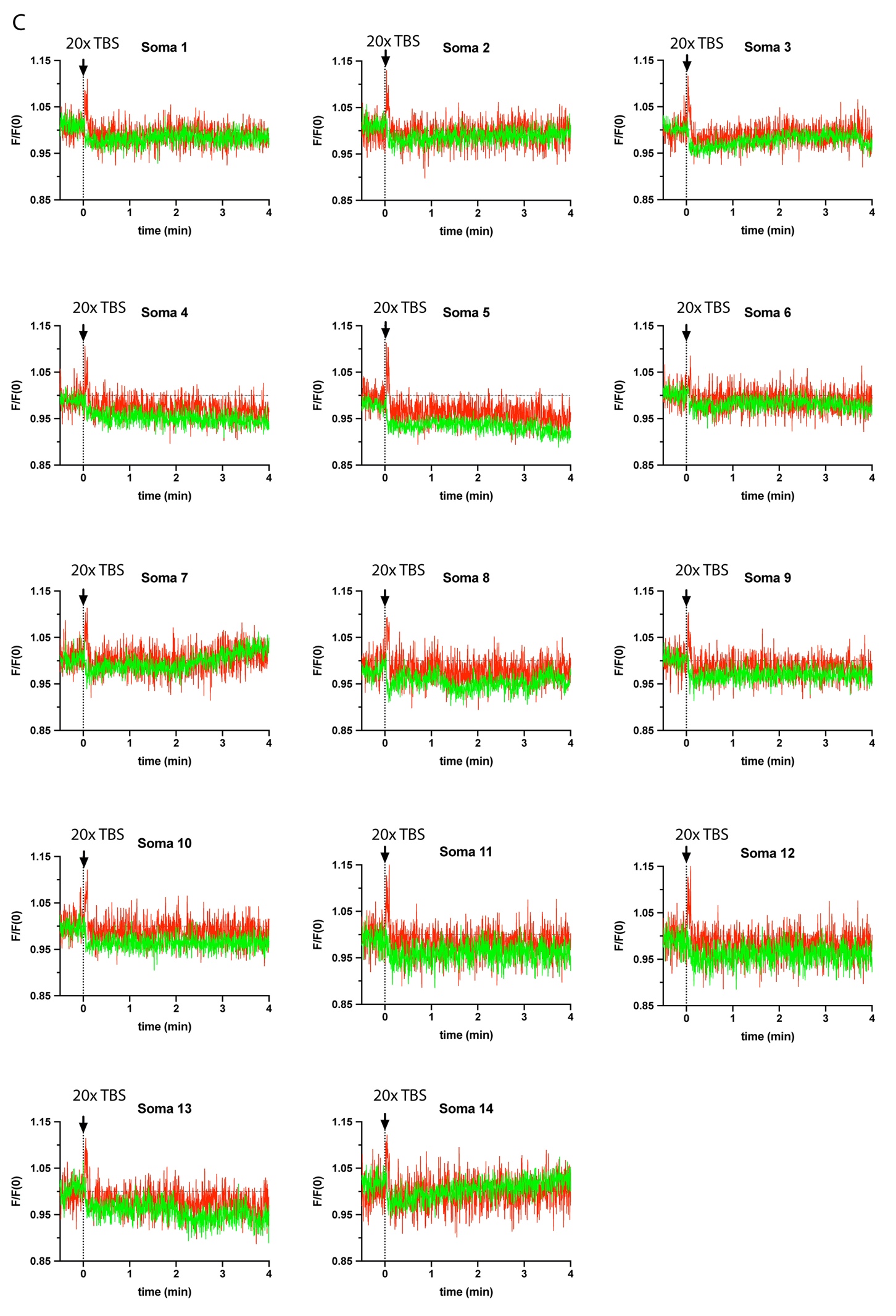

**Slice 3:**

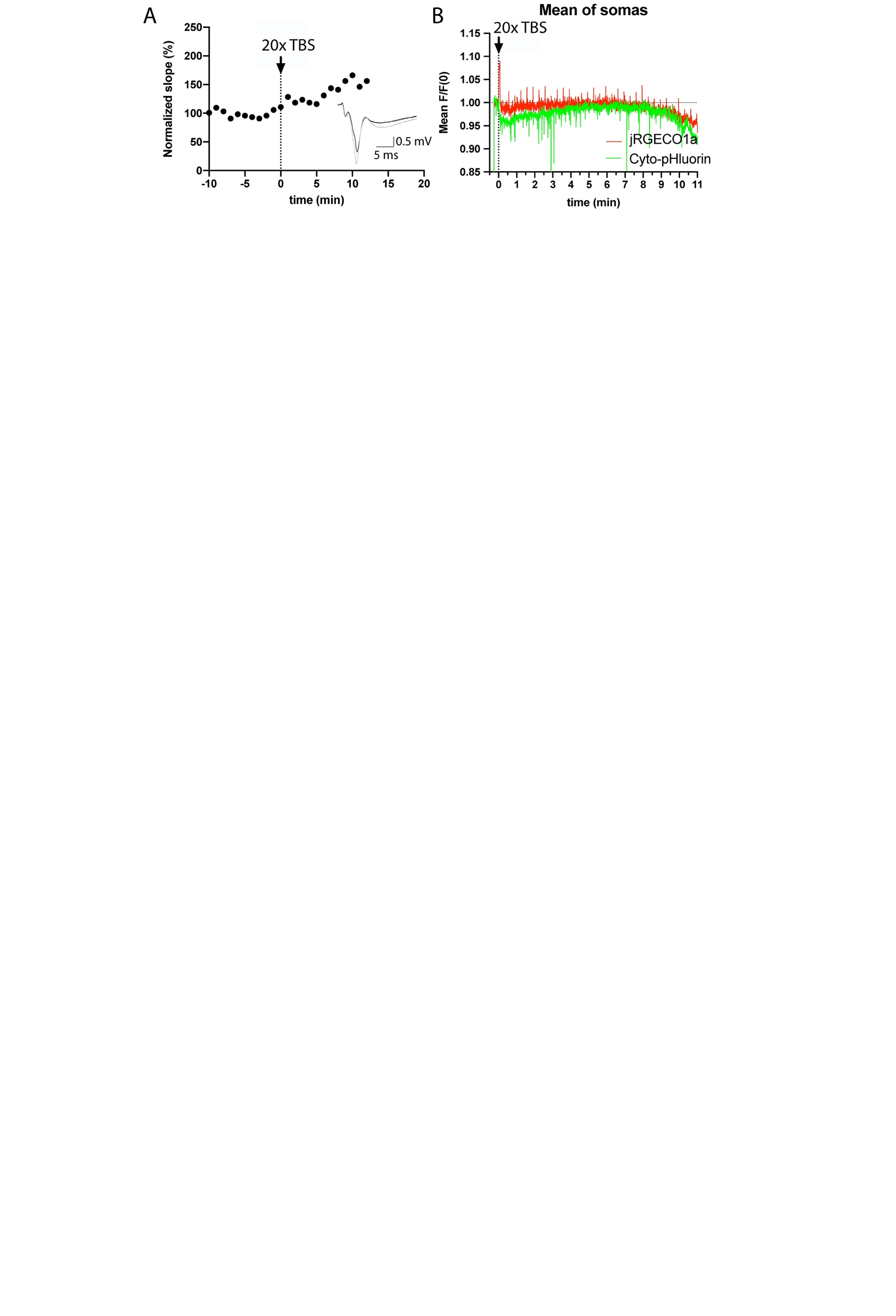

*(“Slice 3” continued on next page)*

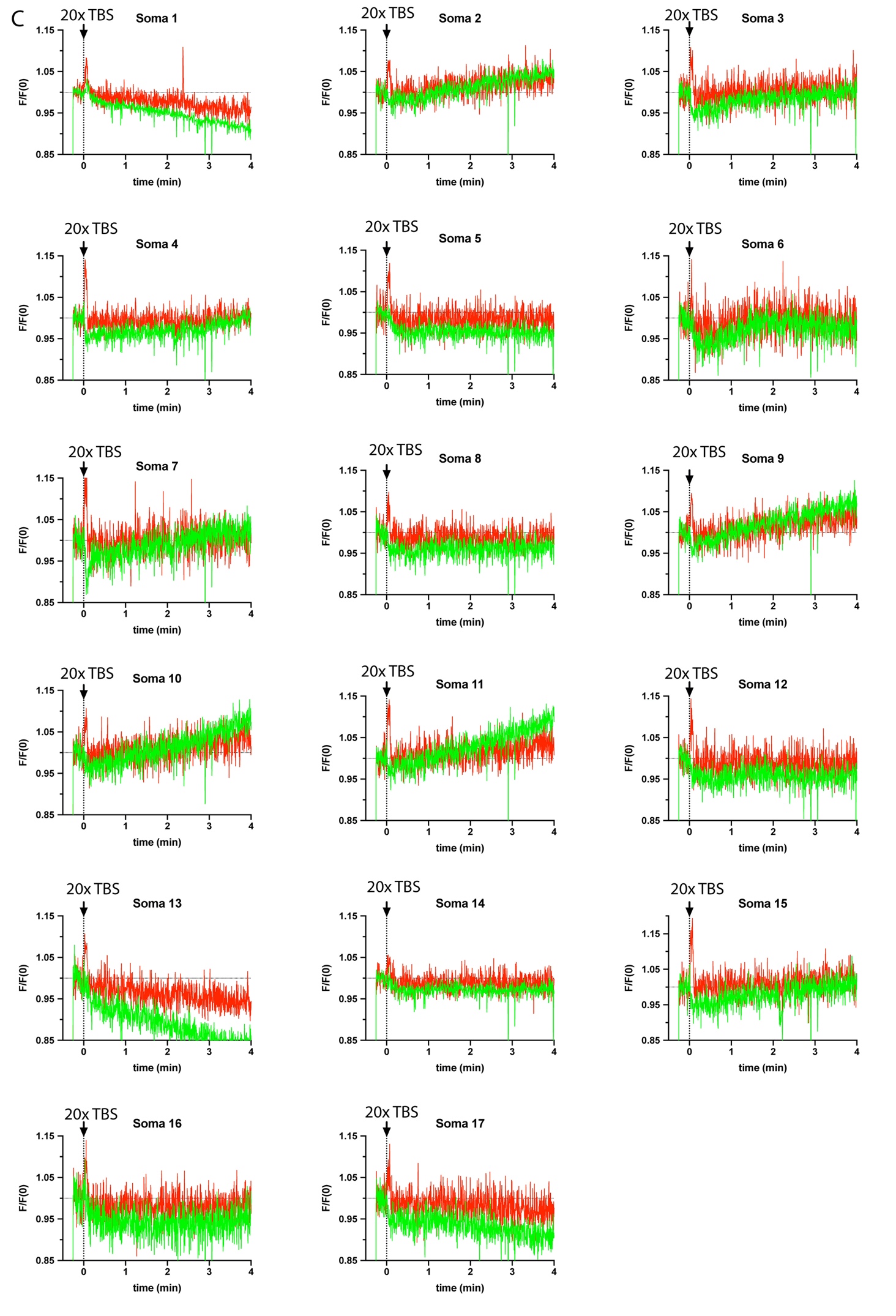
